# Selection for a shared behavior creates parameter dependence read as synergy in populations of neuron models

**DOI:** 10.64898/2026.11.27.769668

**Authors:** Nurit Degani-Katzav, Alon Korngreen

## Abstract

Recent studies found synergy-dominated dependence among the parameters of neuron-model populations selected for a target behavior and asked whether it reflects coordinated regulation of ion channels. We show that selection itself creates synergy of the kind reported, as the standard Gaussian estimator measures it. In eight implementations of six published conductance-based neuron models, we drew independent parameters and kept sets whose 13 electrical features were measurable and close to the published model’s. Relative to same-size unselected samples, selection increased synergy in all seven numerically converged models, including six redrawn at random. At the strictest match, synergy was a median of 16 times (range 4.7 to 40) that of same-size samples that met only the requirement of measurable features. Net O-information, which weighs synergy against redundancy, shifted toward synergy in six of seven models and in a six-parameter gene circuit. Selection strengthened pairwise correlations, which the Gaussian estimator reads as synergy. An exact calculation shows that a single selection constraint makes every parameter triplet synergistic. A nearest-neighbor estimator found that selection also created non-linear pairwise dependence in five of six models. In samples of 50 to 250 models, as in biological datasets, the summary statistic’s sign reversed in 11% to 23% of cases on average, and in up to 48% in the worst case. Synergy in a model population therefore reflects how it was built as well as the neuron and should be compared with a reference drawn from the same prior and selected by the same rule.

## Introduction

Biophysically detailed models are a primary approach to understanding neuronal computation (Almog and Korngreen, 2016; Poirazi and Papoutsi, 2020). They combine the conductance-based description of action potentials (Hodgkin and Huxley, 1952) with compartmental representations of dendritic trees (Rall, 1964), and became practical with dedicated simulators (Hines, 1984; Hines and Carnevale, 1997; Bower and Beeman, 1998). This approach has yielded detailed models of hippocampal, cerebellar, and neocortical neurons (Traub et al., 1991; De Schutter and Bower, 1994; Mainen and Sejnowski, 1996; Hay et al., 2011). Historically, recordings were reduced to best-fit channel parameters, assembled into a single deterministic model, and tuned by hand or automated search to reproduce selected firing features (Vanier and Bower, 1999; Keren et al., 2005; Druckmann et al., 2007; Gouwens et al., 2018). Each step is legitimate, but this one-way pipeline collapses experimental variability and fitting uncertainty into a single parameter set (Korngreen, 2026a, 2026b).

Research on the crustacean stomatogastric ganglion revealed what this compression obscures. Simulations of more than 20 million model circuits showed that virtually indistinguishable network activity can arise from widely different synaptic strengths and intrinsic properties (Prinz, Bucher and Marder, 2004). Model neurons with similar firing can differ sevenfold in a single conductance (Goldman et al., 2001), and models built from the average conductances of similar models may fail to reproduce their behavior (Golowasch et al., 2002). Neurons of the same type also vary severalfold in channel expression and maximal conductances (Schulz et al., 2007; Marder and Taylor, 2011). Degeneracy is therefore a property of biological systems, not merely of their models (Edelman and Gally, 2001; Marder and Goaillard, 2006; Goaillard and Marder, 2021), with parallels in hydrology (Beven, 2006), statistics (Breiman, 2001), metabolic modeling (Reed and Palsson, 2004), and machine learning (D’Amour et al., 2022). If many parameter sets fit the data equally well, the scientific object is an ensemble of models, not a single best parameter set (Marder and Taylor, 2011; Korngreen, 2026b). Such populations have been built by sampling databases of parameter sets (Prinz et al., 2003), mapping the solutions of automated searches (Achard and De Schutter, 2006), multi-objective optimization (Druckmann et al., 2007), calibration to experimental variability (Britton et al., 2013), and simulation-based inference, which returns a posterior distribution over parameters (Gonçalves et al., 2020). Even the Hodgkin–Huxley model behaves as such a population: its excitability is set by a few parameter combinations (Ori et al., 2018), and reintroducing the uncertainty of the original rate-constant fits produced a heterogeneous population in which the canonical 1952 parameter set was a minority (Korngreen, 2026a).

Ensemble thinking, however, relocates the methodological problem instead of resolving it. A model population is also a statistical object produced by a procedure: a prior from which parameters are drawn, protocols and features that evaluate them, and a rule that decides which parameter sets are retained. We recently argued that reusable neuron models should carry a declared scope and an explicit account of their uncertainty, so that a conclusion can be traced either to the data or to a modeling choice (Korngreen, 2026b).

Pairwise analyses of populations of acceptable models often found weak parameter correlations (Taylor et al., 2009; Britton et al., 2013; Rathour and Narayanan, 2014), prompting research into higher-order dependencies. The O-information (Rosas et al., 2019) summarizes how a group of variables shares information: positive values indicate redundancy and negative values synergy, following a decomposition of multivariate information into these components (Williams and Beer, 2010). Applying this measure to more than 110,000 layer 5 pyramidal neuron models, Reva and colleagues (2025) found synergy roughly ten times stronger than redundancy, with the most synergistic groups comprising 25 parameters. Gene-expression measurements in single dopaminergic neurons likewise suggest that up to 41% of information shared by co-varying ion-channel and dopamine-metabolism genes lies beyond pairwise relationships (Tapia et al., 2018). Reva and colleagues proposed such stochastic model populations as a null model for biological variability, against which departures might reflect transcriptional co-regulation or methodological factors such as the sampling strategy, and read the synergy as functional coordination rather than a co-regulated gene module (Reva et al., 2025).

In a population of models, however, O-information summarizes the dependence among parameter values after the population has been constructed. It cannot tell whether the dependence came from the prior, the acceptance rule, or a biological regulatory process, nor does it establish dynamical interactions in any single model. Moreover, selection for a shared behavior can create dependence among parameters whenever the acceptance rule couples them, whatever the biology. Keeping a parameter set because its output matches a target is a form of conditioning on a common effect: multiple parameters can contribute to the same output, so conditioning on that output can induce dependence among parameters that were independent under the prior (Pearl, 1995). Epidemiology has documented how large such selection effects can be (Cole et al., 2010; Hernán et al., 2011; Munafò et al., 2018; Griffith et al., 2020). In models, requiring a function to hold creates negative correlations among independently generated variables (Schneider et al., 2023), and confining model inputs to a constrained region makes them dependent and changes how output variance is attributed among them (Kucherenko et al., 2017). In neuron modeling, Arnaudon and colleagues (2023) traced a correlation between somatic and axonal sodium conductance partly to the constraint on spike amplitude, and constraining a compartmental model only with somatic recordings, or with reduced features of the dendritic membrane potential, yields a large solution manifold (Keren et al., 2005; Almog and Korngreen, 2014).

Two further problems complicate the interpretation of higher-order statistics in model populations. The first is the size of the search. The number of groups grows combinatorially with the number of parameters and with group size (Neri et al., 2024): a model with 30 parameters has 4,060 triplets and 27,405 groups of four. The most extreme groups must therefore be judged against what an equally large search of unstructured data would find (Benjamini and Yekutieli, 2001). The second is the estimator. The O-information is usually computed assuming that the parameters are jointly Gaussian, in which case it depends only on their covariance matrix (Ince et al., 2017); for weak correlations, its value for three parameters is approximately the product of their three pairwise correlations (Methods), so this measure of higher-order dependence is a function of pairwise statistics. Even simple Gaussian systems often carry net synergy (Barrett, 2015), and in small systems the O-information can report net redundancy where synergy dominates (Varley et al., 2023).

This issue is distinct from whether channels are co-regulated. Genuine co-regulation certainly exists. Channel genes are co-expressed in cell-type-specific patterns in identified crustacean neurons (Schulz et al., 2007); these correlations are maintained by neuromodulatory input, not synaptic activity (Temporal et al., 2012), and extend to neurofilament and synaptic genes (Kodama et al., 2020). O’Leary and colleagues (2013) showed that a homeostatic tuning rule produces correlations among conductances like those measured in identified neurons, which populations of models selected from random parameter sets for the same activity had not shown (Taylor et al., 2009). Biological co-regulation and dependence created by selection can therefore coexist in the same population. The question is whether a statistic computed on a selected population of models can tell them apart.

Here we ask a narrower question: how much synergy, as the usual estimator measures it, does selection create when parameters are sampled independently, and how much remains after controlling for search size and population size? Such dependence is routinely measured in model populations selected for a target behavior and attributed to the modeled neuron, but the contribution of selection alone has not been measured. We also ask whether common estimators detect non-linear dependence, whether selection creates dependence beyond pairwise correlations, and whether the balance of synergy and redundancy reflects population construction rather than the modeled neuron.

We hypothesized that selecting parameter sets by their electrical output is sufficient to produce synergy-dominated dependence, as measured by the Gaussian estimator, even when the parameters are drawn independently (a factorized prior) and are not co-regulated. Because selection operates through the model’s mapping from parameters to features, the resulting dependence reflects both the model and the acceptance rule. To test this, we drew 38,400 parameter sets from a factorized prior with an evenly spread quasi-random (Sobol) design for eight implementations, or code versions, of six published conductance-based neuron models, simulated them and selected populations at four levels of stringency. We repeated the key comparisons on parameter sets drawn at random for six of the implementations. We measured higher-order dependence against unselected samples of equal size and controls for search size, estimator, and population construction.

We report three findings. First, selection alone made the parameters synergistic as the Gaussian estimator measures it: relative to unselected samples of the same size, the size of the synergy rose in every converged model, and at the strictest tolerance it was a median of 16 to 19 times that of same-size samples that met only the requirement that all features be measurable. Net O-information shifted toward synergy in six of seven models, and the same selection shifted the six-parameter O-information of a gene-circuit model toward synergy. Second, the effect lies in the strength of the correlations created by selection, which the Gaussian estimator reads as synergy, as an exact calculation for a single constraint shows; selection also created non-linear pairwise dependence in five of six models, whereas the average over parameter triplets showed no consistent dependence beyond what the correlations describe. Third, identifying synergistic parameter groups depended on population size: groups passing a search-size control were rare in the two smallest populations, and the sign of the summary statistic was unreliable at the sizes of biological datasets. The internal statistical structure of an ensemble therefore cannot be interpreted apart from the procedure that produced it.

## Methods

### Model panel

We used eight implementations of six published neuron models (Table 1; Supplementary Table S1 summarizes all numerical and analysis settings). Because published models and channel mechanisms are widely reused (Almog and Korngreen, 2016), we distinguish a source model, the original published model, from an implementation, one code version of it. Implementations of one source model share equations and parameter conventions and are not independent tests, so summaries across models were also computed with the source model as the unit. Below, “model” means an implementation unless stated otherwise. The panel contained two Hodgkin and Huxley (1952) implementations (one compartment, and a soma with a ten-compartment dendrite), the regular-spiking and fast-spiking cortical models of Pospischil and colleagues (2008), the cerebellar Purkinje model of Khaliq and colleagues (2003), the hippocampal CA1 pyramidal model of Migliore and colleagues (1999), and two layer 5 cortical pyramidal models (Mainen and Sejnowski, 1996; Hay et al., 2011). The Mainen model failed the convergence test described below: refining the mesh lost a near-threshold spike and changed the adaptation index in one of seven test parameter sets. We excluded it from all summaries but show it in the figures; including it changed no qualitative conclusion (Supplementary Section S1). The primary panel therefore had seven converged implementations of five source models. Two Hay-model parameters (the apical Kv3-like potassium and low-voltage-activated calcium conductances) changed no output when scaled from one quarter to four times their value and were removed before analysis, because they add only noise to the dependence measures (Supplementary Section S1). No neuron class is represented by two independent source models.

**Table 1.**
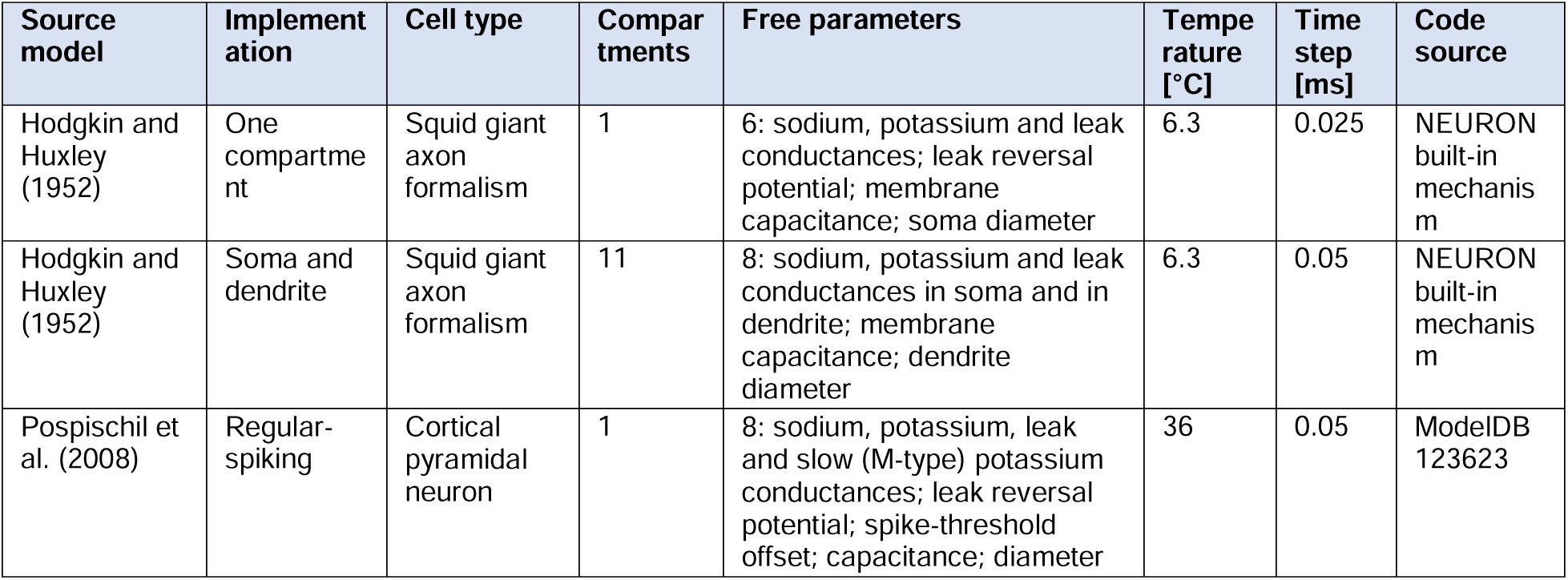
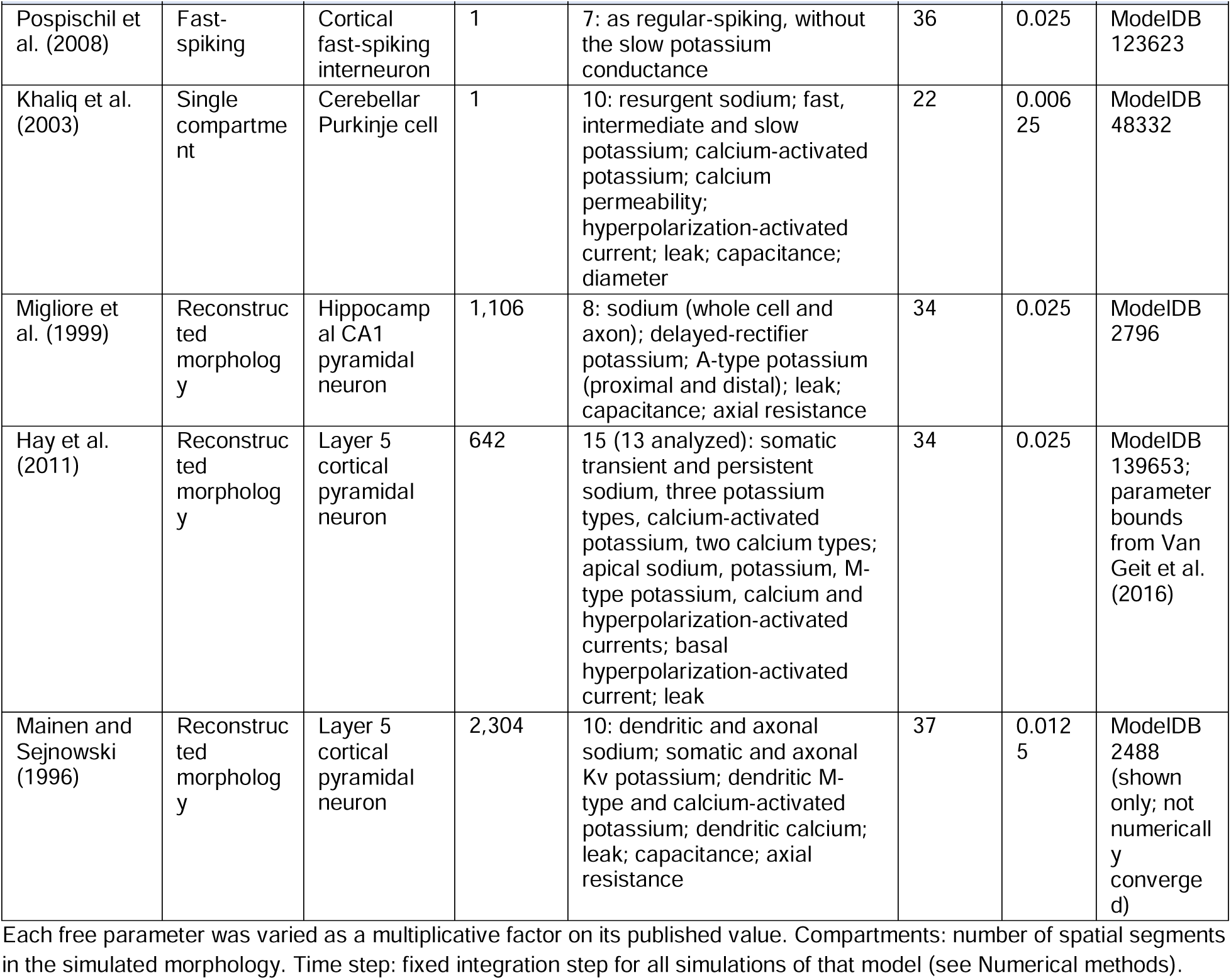
Neuron models used in this study.

| Source model | Implementation | Cell type | Compartments | Free parameters | Temperature [°C] | Time step [ms] | Code source |
|---|---|---|---|---|---|---|---|
| Hodgkin and Huxley (1952) | One compartment | Squid giant axon formalism | 1 | 6: sodium, potassium and leak conductances; leak reversal potential; membrane capacitance; soma diameter | 6.3 | 0.025 | NEURON built-in mechanism |
| Hodgkin and Huxley (1952) | Soma and dendrite | Squid giant axon formalism | 11 | 8: sodium, potassium and leak conductances in soma and in dendrite; membrane capacitance; dendrite diameter | 6.3 | 0.05 | NEURON built-in mechanism |
| Pospischil et al. (2008) | Regular-spiking | Cortical pyramidal neuron | 1 | 8: sodium, potassium, leak and slow (M-type) potassium conductances; leak reversal potential; spike-threshold offset; capacitance; diameter | 36 | 0.05 | ModelDB 123623 |
| Pospischil et al. (2008) | Fast-spiking | Cortical fast-spiking interneuron | 1 | 7: as regular-spiking, without the slow potassium conductance | 36 | 0.025 | ModelDB 123623 |
| Khaliq et al. (2003) | Single compartment | Cerebellar Purkinje cell | 1 | 10: resurgent sodium; fast, intermediate and slow potassium; calcium-activated potassium; calcium permeability; hyperpolarization-activated current; leak; capacitance; diameter | 22 | 0.00625 | ModelDB 48332 |
| Migliore et al. (1999) | Reconstructed morphology | Hippocampal CA1 pyramidal neuron | 1,106 | 8: sodium (whole cell and axon); delayed-rectifier potassium; A-type potassium (proximal and distal); leak; capacitance; axial resistance | 34 | 0.025 | ModelDB 2796 |
| Hay et al. (2011) | Reconstructed morphology | Layer 5 cortical pyramidal neuron | 642 | 15 (13 analyzed): somatic transient and persistent sodium, three potassium types, calcium-activated potassium, two calcium types; apical sodium, potassium, M-type potassium, calcium and hyperpolarization-activated currents; basal hyperpolarization-activated current; leak | 34 | 0.025 | ModelDB 139653; parameter bounds from Van Geit et al. (2016) |
| Mainen and Sejnowski (1996) | Reconstructed morphology | Layer 5 cortical pyramidal neuron | 2,304 | 10: dendritic and axonal sodium; somatic and axonal Kv potassium; dendritic M-type and calcium-activated potassium; dendritic calcium; leak; capacitance; axial resistance | 37 | 0.0125 | ModelDB 2488 (shown only; not numerically converged) |

Each free parameter was varied as a multiplicative factor on its published value. Compartments: number of spatial segments in the simulated morphology. Time step: fixed integration step for all simulations of that model (see Numerical methods).

### Parameter sampling

Each parameter was a multiplicative factor on its published value (published model = 1), drawn independently for each parameter and uniformly on a logarithmic scale (a factorized log-uniform prior), with a scrambled Sobol design, an evenly spread quasi-random sequence (Sobol’, 1967). Ranges were one quarter to four times the published value for conductances, narrower for capacitance, axial resistance, reversal potentials and morphology, and, for the Hay model, the bounds of its published optimization (Van Geit et al., 2016). These are expert-chosen or optimization ranges, not measured biological distributions, so no result below describes biological variability. Sample size was set by computational cost: 8,192 parameter sets for each of the two Hodgkin–Huxley and two Pospischil implementations, 4,096 for the Khaliq model, 1,024 for the Hay model and 256 each for the Migliore and Mainen models, 38,400 in total (the unselected sample). Sets whose membrane potential left −150 to +80 mV were removed as numerical failures; this occurred only in the Khaliq model, where 3,501 of 4,096 sets remained. Because the prior contains no relationship between parameters, any later dependence was introduced by operations on the sample. A finite Sobol design, however, spreads points more evenly than independent random draws and has smaller sample correlations; we controlled for this with unselected samples and shuffled copies of the same size. To test whether the results depend on this design, we redrew the prior with independent pseudo-random draws and repeated the selection for six implementations: the five with the largest selected populations (the two Hodgkin–Huxley, the two Pospischil and the Khaliq implementations), at the same sample sizes (3,525 of the 4,096 Khaliq sets passed the voltage check), and the Hay model, with 8,192 sets because its 1,024 were too few to resolve the effect. This random-draw run added 45,056 parameter sets, for 121,344 simulated in all.

### Stimulation protocols and electrical features

Each parameter set was simulated with common somatic current-clamp protocols: 300 ms without stimulation (resting potential) and, after a 120 ms delay, three 400 ms steps, one hyperpolarizing and two depolarizing, with amplitudes fixed for each model from the published model’s current-to-firing-rate curve. Rheobase, the smallest step current evoking a spike, was found by scanning 64 equally spaced amplitudes and refining the first spiking step by bisection, with a spike defined as an upward crossing of 0 mV during a 140 ms step (Supplementary Section S1). We measured 13 features: resting potential; input resistance, membrane time constant and voltage sag during the hyperpolarizing step; rheobase; and, during the stronger depolarizing step, first-spike latency, spike amplitude, spike width at −20 mV, after-hyperpolarization depth, first interspike interval, mean firing rate, adaptation index and spike count. Spike amplitude, spike width and after-hyperpolarization depth were extracted with eFEL (Mandge et al., 2026); spike count, firing rate, first-spike latency, first interspike interval and adaptation index were computed from upward crossings of −20 mV; all were checked against an independent implementation. The spike width is the width at the −20 mV detection threshold, which in the published parameter sets was 0.1 to 0.3 ms wider than the width at half amplitude (the stored variable carries the name of the latter). A second protocol set, with steps scaled to each parameter set’s rheobase, used 12 features: the same without sag, with the spike features measured at three times rheobase. It is undefined when the rheobase is zero, as for the published Khaliq model, which fires spontaneously, so the Khaliq model enters only the first set. The second set gave qualitatively similar results and enters only summaries stated to average over both protocol sets (Figs. 4A, 4C and 5C, D).

### Selecting populations for a target behavior

The target was the published model’s own feature vector, a deterministic simulation output, not an experimental measurement. Each feature’s distance from the target was expressed in standard deviations of that feature across the whole unselected sample, a scaling computed once, before selection. The 13 distances were combined as the largest single distance or as the root-mean-square distance (the weighted Euclidean distance, every feature weighted 1/13), and a parameter set was accepted if this distance was below 3, 2, 1, or 0.5 standard deviations (Fig. 1A). The largest-distance criterion requires every feature to lie within the tolerance, whereas the root-mean-square criterion lets a large deviation in one feature be offset by small deviations in others and so accepts more sets. Sets with any undefined feature were rejected, not imputed, so acceptance has two stages: all 13 features must be measurable, which requires at least three spikes at the stronger step (for the adaptation index) and the successful measurement of every other feature, including a fit of the membrane time constant; and the distance must lie below the tolerance. In the Migliore model the resting potential did not vary across the unselected sample, so its standardized distance was undefined and omitted. The root-mean-square denominator remained 13, which makes the tolerance about 4% looser; with a denominator of 12, one to eight fewer sets were accepted and the synergy part (defined below) was 5% to 25% larger. Because each step lasts 400 ms, mean firing rate is exactly 2.5 times spike count, so firing output contributes two of the 13 terms of the root-mean-square distance (two of 12 in the second protocol set); we repeated the selection with the rate removed. The accepted sets form a selected population, whose distribution is conditional on the prior, the simulated features, and the acceptance rule; populations with fewer than 60 sets were not analyzed. Acceptance rates are shown in Fig. 3D.

**Fig. 1.**
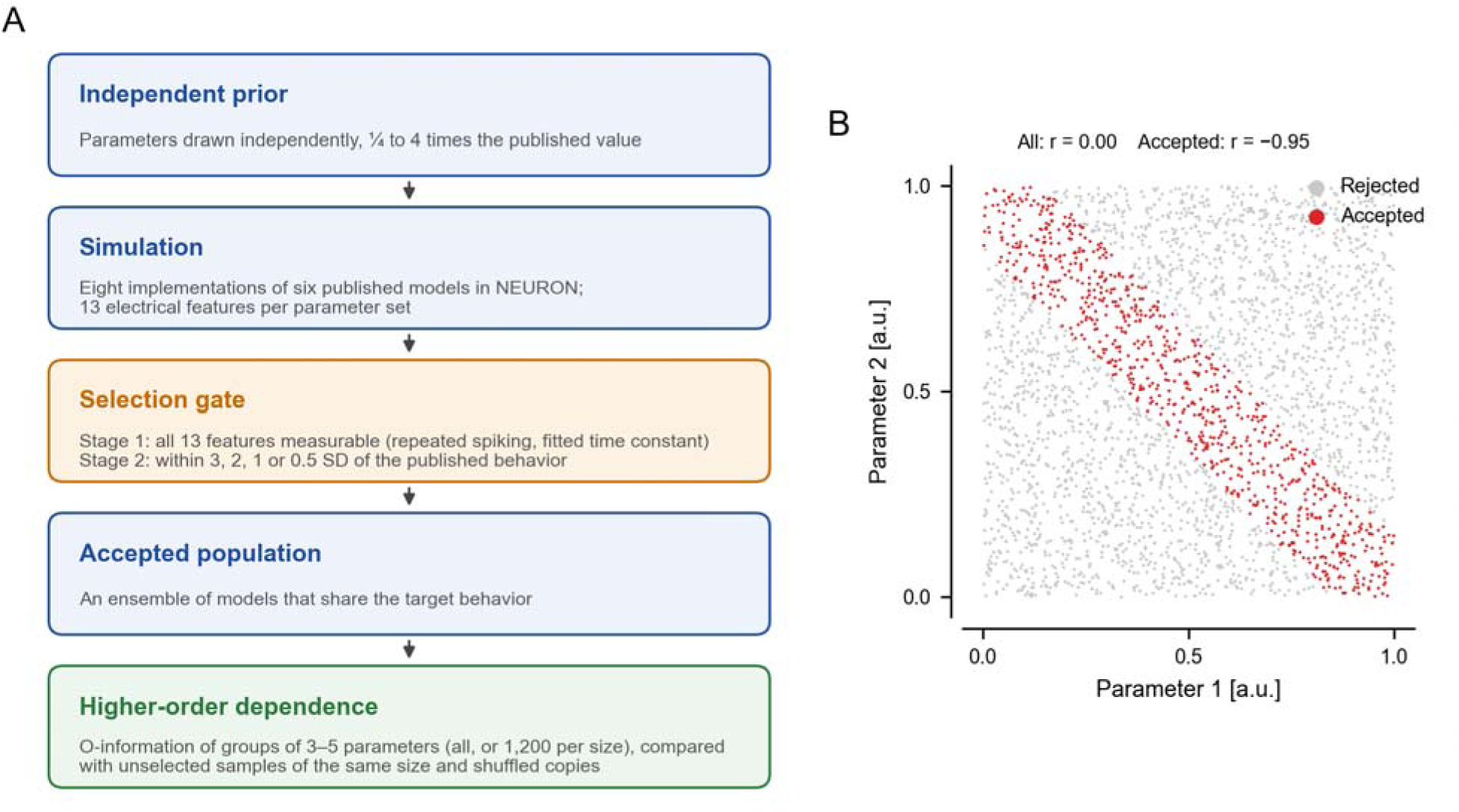
Study design. A, The pipeline used for every model. Parameter sets were generated from a broad factorized prior around each published model (¼ to 4 times the published value for conductances), simulated, and described by 13 electrical features. A two-stage selection gate kept the sets whose 13 features could all be measured and lay within a tolerance of the published model’s features, in standard deviations (SD) of the unselected sample. Higher-order dependence was measured in groups of 3, 4 and 5 parameters of the accepted population (all groups, or 1,200 of each size) and compared with unselected samples of the same size, shuffled copies and a search-matched control. B, Illustration only: two artificial parameters drawn independently and uniformly (3,000 points). Keeping only the points whose sum is within 0.15 of 1 (red; rejected points in gray) creates a strong negative correlation. This pairwise effect is not a measurement of the three-parameter dependence analyzed in the rest of the paper. *r*, Pearson correlation coefficient; a.u., arbitrary units.

### Other ways of building a population

To separate the effect of selection from that of population structure, we also built populations in other ways. Two required additional simulations for the six implementations with at least 1,024 unselected parameter sets, each with half as many sets as the unselected sample (18,944 sets each; 76,288 parameter sets simulated in all): (1) a prior with correlated neighboring parameters (Gaussian copula; correlation 0.6 raised to the power of their distance in parameter order); and (2) a clustered population, sampled with Gaussian jitter (standard deviation 6% of each parameter’s range) around eight parameter sets from the unselected sample; jitter beyond a prior bound was set to the bound, so on average 2.8% to 6.6% of a model’s parameter values are repeats. The others were derived from the population selected with the largest-single-distance criterion at 3 standard deviations: (3) the half lying most strongly along its main covariance directions, chosen within deciles of distance to the target to hold fitness constant; (4) a random subset of the same size as (3); (5) a shuffled copy, with each parameter permuted independently, which keeps every parameter’s distribution but destroys all dependence; and (6) a synthetic population drawn from a Gaussian copula fitted to the normal-score correlations and the distribution of each parameter, without any neuron model. The high-covariance and random subsets were matched in size and parameter means but not in the full shape of each parameter’s distribution (Supplementary Section S1).

### Measuring higher-order dependence

For a group of *n* parameters *X* = (*X*_1_, …, *X*_n_), the O-information is Ω = (*n* − 2) *H*(*X*) + Σ_i_ [*H*(*X*_i_) − *H*(*X*_−i_)], where *H* is entropy, and *X*_−i_ is the group without parameter *i* (Rosas et al., 2019). Positive Ω indicates redundancy and negative Ω synergy; following the literature we address, we call a group synergy-dominated when Ω < 0, with no magnitude threshold. Units are nats (information measured with natural logarithms; 1 millinat = 0.001 nats). We estimated Ω assuming that the logarithms of the parameter factors are jointly Gaussian, so that Ω depends only on the correlation matrix, with an analytical small-sample correction, and with a Gaussian copula, which first converts each parameter to normal scores by rank (tied values receive their average rank) and is therefore unaffected by monotone rescaling (Ince et al., 2017). The two agreed (correlation 0.91 across the 14,291 triplet estimates of the 164 populations of the primary panel, and 0.96 between their shares of synergy-dominated triplets; Supplementary Section S1), and the Gaussian values are reported.

Because a Gaussian estimate is determined by the correlation matrix, a negative value summarizes pairwise structure and does not prove dependence beyond second-order statistics. For population correlations a, b and c, the Gaussian O-information of a triplet is ½ ln[(1 − a² − b² − c² + 2abc) / ((1 − a²)(1 − b²)(1 − c²))], whose leading term for weak correlations is the product abc. A negative product therefore implies synergy, but a positive product does not guarantee redundancy. If independent, standardized Gaussian parameters are selected for a narrow band of one weighted sum, with unit weight vector u, the variance along u shrinks by a fraction k, the covariance becomes I − k uu^T^, and every triplet is synergistic, with O-information approximately −k^3^u_a_^2^u_b_^2^u_c_^2^, the product of its three correlations (Supplementary Section S1). Across random correlation matrices with a mean absolute correlation of 0.04, as in the selected populations, the exact value and the product correlated at 0.99 and had the same sign 98% of the time; in the estimates, which include the small-sample correction, the signs agreed for 285 of 287 triplets in the five large populations selected at 3 standard deviations but for only 44 of 56 and 225 of 286 in the Migliore and Hay populations (154 and 344 sets). Ω was computed for every group of 3, 4 and 5 parameters, or for a fixed random sample of 1,200 groups when there were more. Each population was summarized by the share of synergy-dominated triplets and by the net O-information, Ω_net_ = (1/3) Σ_n=3_^5^ (1/*n*) [ max(Ω, 0) _n_ − max(−Ω, 0) _n_], where · _n_ is the mean over the analyzed groups of size *n*. The first term is the redundancy part of the net O-information and the second, the mean size of the negative values, the synergy part; we also report both separately. Negative net O-information indicates synergy overall. The share counts only signs, whereas the net O-information also weighs their size. Pairwise dependence was summarized as the mean absolute Pearson correlation between the logarithms of the parameter factors over all parameter pairs. Throughout, “synergy” is a statistical property of a population of parameter sets; it does not imply that the parameters interact in the dynamics of the model, which requires perturbing the model directly.

### Comparison with unselected samples of the same size

Every statistic used here depends on the number of parameter sets, and the selected populations were smaller than the unselected sample, whose quasi-random design also has unusually small sample correlations (Supplementary Section S1). The full unselected sample is therefore not a fair reference for a selected population. We compared each selected population with 100 random subsamples of the unselected sample of exactly the same size, analyzed identically, and called a selected value different from chance if it lay outside their central 95% (the chance range). We also compared it with 20 shuffled copies of itself, which keep each parameter’s distribution but contain no dependence and behave like independent random draws. The first comparison controls population size within the actual sampling design; the second approximates a null of independent random samples. To separate the two stages of acceptance, we also compared each selected population with 100 samples of the same size drawn from the unselected parameter sets whose 13 features are all measurable (feature-complete samples). Both comparisons were repeated on the parameter sets drawn at random. The values for each model are listed in Supplementary Section S2.

### Control for the size of the search

To assess whether an individual group would appear unusually synergistic in a search of the same size over data without cross-parameter dependence, we built 25 shuffled copies of each population, computed Ω for the same number of groups of each size, and recorded the smallest and largest value in each copy. The 5th percentile of the minima and the 95th percentile of the maxima defined an exploratory search-matched surrogate band (Theiler et al., 1992; Schreiber and Schmitz, 1996); a group with Ω below its lower bound was described as falling beyond this surrogate threshold. With only 25 shuffled copies, the tail thresholds are imprecise, and the band is not a validated family-wise error-rate procedure. Because shuffling also destroys dependence introduced by selection, this comparison does not control for selection-induced dependence.

### Which features determine selection?

To identify the features through which selection acts, we rebuilt every population selected at 3 and 2 standard deviations with each of the 13 features left out of the distance in turn. For each omission we recorded two things. The first was the share of accepted parameter sets that changed, split into sets that became eligible because the left-out feature had been undefined for them and sets whose acceptance changed through the distance alone (which, with the root-mean-square criterion, then averages over 12 features). The second was the change in the synergy part of the net O-information, both for the new population and with the feature-complete population held fixed. Leaving out rheobase tests its contribution to these acceptance rules at 3 and 2 standard deviations; it does not show what a differently measured rheobase would do (Discussion, Limitations). To ask whether the same groups were identified at different tolerances, we compared the (up to) ten most synergistic triplets, those with negative O-information, at each tolerance with those at 3 standard deviations, and with those of 200 random subsets of the 3-standard-deviation population of the same size. The results for each feature and model are listed in Supplementary Section S2.

### Estimator test systems

We tested the estimators on eleven artificial systems of known dependence. Seven had four variables. One had none (four independent variables; expected Ω = 0). Three were redundant (expected Ω > 0): linearly correlated variables, three noisy copies of one variable, and a pooled mixture of two populations with different means. Three were synergistic (expected Ω < 0): the fourth variable equaled, up to a small noise, the sum of the other three, the product of two of them, or the product of all three. The product systems used zero-mean variables, so that every pairwise linear correlation vanishes; the variables remain dependent, jointly and in pairs through the spread of the product, so these systems test dependence that the correlation matrix does not capture. The product of two such variables is the continuous analog of an exclusive-or relation. Four systems represented selection on positive parameters, drawn log-uniformly from one quarter to four times a reference value and analyzed as logarithms: the product of three such variables, and three selection bands, in which three parameters were drawn and only the 10% whose score lay closest to its median were kept, the score being the sum of the log-parameters (a multiplicative constraint), the sum of their squares or x_1_ + |x_2_ − x_3_| (both non-monotone). These four were sampled with five random seeds at each size and averaged. Each system was sampled at 300, 1,000, 4,000 and 16,000 points and analyzed with the Gaussian and Gaussian-copula estimators and with a nearest-neighbor entropy estimator that makes no distributional assumption (Kozachenko and Leonenko, 1987; fourth nearest neighbor, maximum-coordinate distance). Estimates above +0.02 nats were classified as redundant, below −0.02 nats as synergistic and otherwise as no dependence, and scored against the known dependence. The Gaussian estimators need only a covariance matrix; the nearest-neighbor estimator searches the neighbors of every sample, so its cost grows with sample size, and it needs several thousand samples to be reliable (Results).

### Nearest-neighbor check on selected populations

To test for dependence that the Gaussian estimator cannot see, we applied the nearest-neighbor estimator to every parameter triplet and parameter pair of the selected populations drawn at random that contained at least 2,000 parameter sets (27 populations from six models, 2,002 to 7,101 sets; both distance criteria; populations at 3, 2 and 1 standard deviations qualified, none at 0.5). Each parameter was first converted to normal scores by rank, so that the estimate reflects only the dependence between parameters. Each population was compared with ten of each of three references: unselected samples of the same size; shuffled copies; and synthetic populations with the same rank correlations and the same distribution of each parameter (Gaussian copula), which keep the dependence that the correlations describe and remove anything beyond it. Because the synthetic populations match the correlations but not the full joint distribution of each pair, the difference from them reflects any dependence the correlations miss, within pairs or among three parameters; it is not a test of three-way dependence alone, and its mean over triplets can hide departures of opposite sign. The excess of pairwise information over the synthetic populations was calibrated against 10 unselected samples of the same size, each compared with five synthetic populations of its own, and checked with a second estimator of mutual information (Kraskov et al., 2004; 39 synthetic populations). Gaussian estimates were also recomputed on the normal scores used by the nearest-neighbor estimator. Parameter sets drawn at random were needed because this estimator responds to the spacing of points: a selected population keeps every point of the evenly spread Sobol design inside a region, whereas random subsamples, shuffled copies and synthetic populations do not (Supplementary Section S2). The same comparisons were applied to the high-covariance and random subsets of these populations and to the correlated prior and clustered population, which were drawn at random. Smaller populations were not analyzed, because this estimator needs several thousand samples (Fig. 2B).

**Fig. 2.**
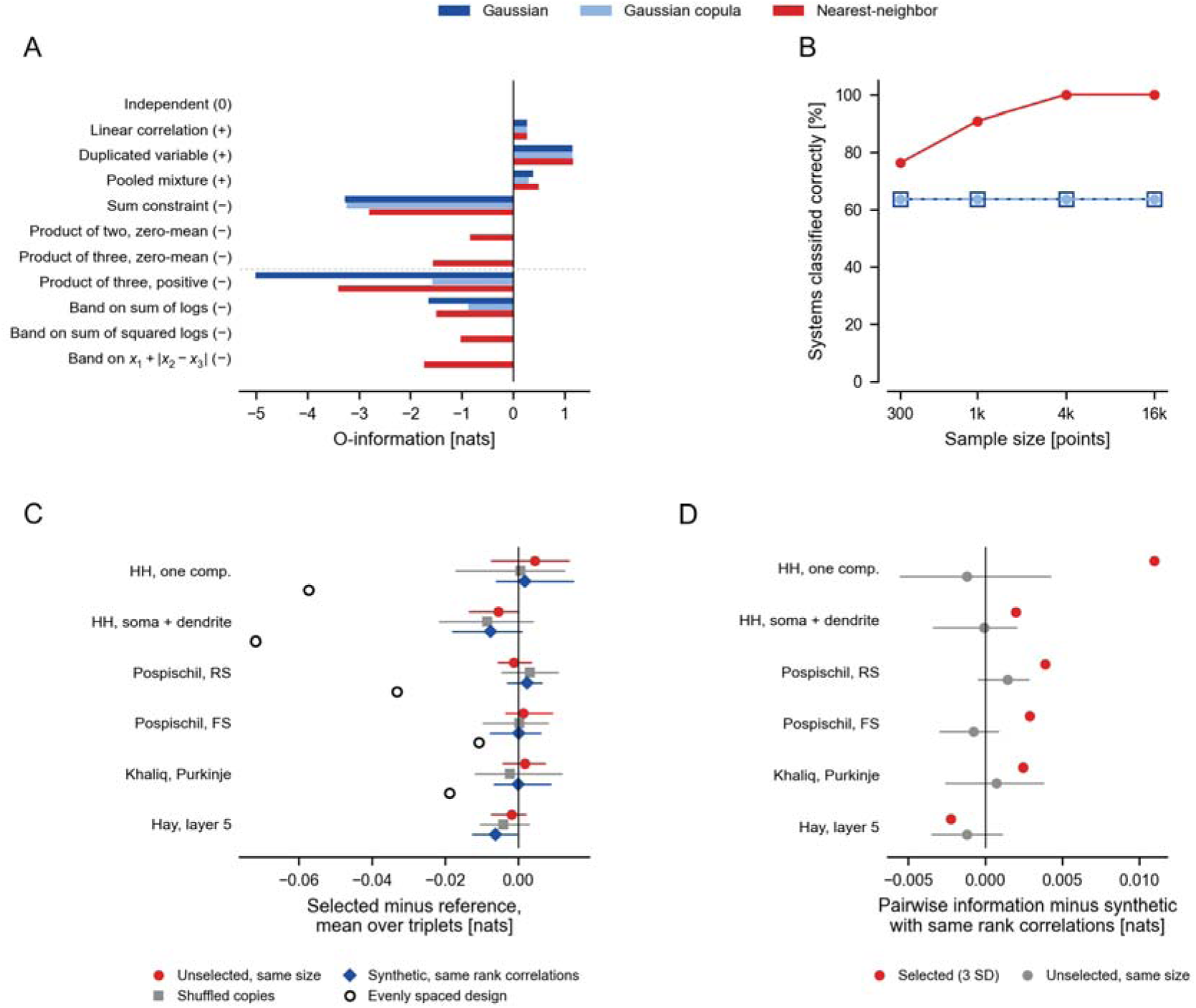
The Gaussian estimator detects some selection constraints and misses others, and selection creates pairwise dependence that the correlations miss. A, O-information of eleven artificial systems of known dependence (16,000 points each) from the Gaussian (dark blue), Gaussian-copula (light blue) and nearest-neighbor (red) estimators; the expected sign is in brackets. Above the dotted line, four-variable systems; in the sum constraint and the two product systems, the fourth variable equals the sum of the other three, the product of two of them or the product of all three (zero-mean variables), plus a small noise. Below it, systems built from parameters drawn log-uniformly and analyzed as logarithms: the product of three positive variables, and three-parameter selection bands that keep the 10% of draws whose score lies closest to its median (means of five seeds). Bars too short to see are within 0.003 nats of zero. B, Percentage of the eleven systems classified correctly (redundant, synergistic or no dependence; threshold ±0.02 nats) as a function of sample size (k, thousand); the Gaussian and Gaussian-copula lines overlap. C, Nearest-neighbor O-information of parameter triplets in the populations selected at 3 SD with the root-mean-square distance from parameter sets drawn at random (2,433 to 7,101 sets), minus that of three references: unselected samples of the same size (red circles), shuffled copies (gray squares) and synthetic populations from a Gaussian copula fitted to the same rank correlations and parameter distributions (blue diamonds). Markers, mean over all triplets and ten references; lines, range over the references; individual triplets can depart in opposite directions; negative values indicate a shift toward synergy. Open circles, the comparison with unselected samples on the evenly spread Sobol design, which is confounded by the spacing of its points (Methods); not available for the Hay model. D, Nearest-neighbor mutual information between parameter pairs minus that of five synthetic populations with the same rank correlations, for the selected populations in C (red) and for 10 unselected samples of the same size, each compared with five synthetic populations of its own (gray; mean and range). Each parameter was converted to normal scores by rank before the nearest-neighbor estimates in C and D. HH, Hodgkin–Huxley; RS, regular-spiking; FS, fast-spiking.

### Reliability of the sign at small population sizes

The biological analyses of higher-order dependence that we discuss used 75 neurons of one transcriptomic type in the mouse visual cortex (Reva et al., 2025) and 111 dopaminergic neurons (Tapia et al., 2018). To measure how reliable the sign of the net O-information is at such sizes, we drew 60 random subsamples of 50, 66, 100, 150 and 250 parameter sets from each of five selected populations whose own sign was stable (largest-single-distance criterion at 1 and 0.5 standard deviations; four implementations of two source models, Hodgkin–Huxley and Pospischil; 291 to 1,243 parameter sets), and recorded the share of subsamples with the opposite sign. Subsamples of 250 from the population of 291 contain nearly all of it, so they vary less than independent samples would.

### Non-neuronal comparison systems

To test whether the effect of selection depends on neuronal dynamics, we applied the same procedure to a six-parameter repressilator, a three-gene oscillating circuit (Elowitz and Leibler, 2000), and a six-parameter three-species Lotka–Volterra community. For each, 20,000 parameter sets were drawn independently and log-uniformly over one decade (repressilator) or 0.8 decades (Lotka–Volterra) around a reference set. Six output features were measured: period, amplitude, mean level, number of peaks, variability of peak intervals and the logarithm of the mean level for the repressilator, so that the mean level enters twice; the three mean abundances, total biomass (their sum), oscillation amplitude and number of surviving species for the community. Populations were selected with the largest-single-distance criterion at the same four tolerances, with the reference set’s features as the target, and the O-information of all six parameters was compared with that of 200 random subsamples of the unselected sample of the same size.

### Attributing variation across models

To ask how much of the variation in synergy across populations was associated with how a population was built and how much with the model it came from, we regressed the synergy part of the net O-information of each population on three non-overlapping groups of predictors: sampling design (construction method, tolerance, logarithm of population size and acceptance rate); population geometry (intrinsic dimension estimated in two ways, relative to the number of parameters); and source model. Geometry measures computed from the covariance matrix (mean absolute correlation, effective rank, participation ratio and the share of variance on the first principal component) were not used, because the synergy part is itself a function of the correlation matrix and would be explained by them partly by construction (Supplementary Section S1). We used ordinary least squares on the standardized outcome, with categorical predictors as fixed indicator variables and no random effects. For each group we report the variance it explains on its own and its unique variance (the R² of the full regression minus that of the regression without the group), which does not depend on how the numerical predictors are scaled; shared variance is not assigned to any group. Intervals were obtained from 2,000 resamples of source models with replacement, skipping the five draws that contained a single source model. Sampling design mixes choices made in advance (construction method and tolerance) with outcomes that depend on the model (acceptance rate and population size). The analysis used 164 populations of the primary panel, one for each combination of model, construction method, tolerance, distance criterion and protocol set (two of them, the Hay populations selected at 3 and 2 standard deviations with fixed-amplitude steps and the root-mean-square criterion, contain the same 344 parameter sets; the two Khaliq populations selected with the undefined second protocol set are excluded). It was repeated without the six clustered populations and for the 93 populations selected for behavior only. Implementation and model architecture were not used as predictors, because each is fixed within a source model and so adds no information beyond it (Supplementary Section S1 and Supplementary Table S2).

### Prediction for an unseen source model

To test whether the relation between construction and synergy carries over to an unseen model, we left out each source model in turn, fitted the regression on sampling design and intrinsic dimension for the others, and computed the R² of the prediction for the left-out model relative to predicting the training models’ mean for each of its populations; a positive value means that the prediction beat this guess. We also scored the same predictions against the left-out model’s own mean, which a prediction beats only if it is accurate in absolute value within that model. All preprocessing (scaling, encoding of categories, imputation) was learned from the training models only. Predictions that also used the covariance-based geometry measures, and predictions from sampling design alone, are reported in Supplementary Section S1.

### Statistical analysis

Agreement in the direction of an effect across source models was tested with an exact two-sided sign test. With five source models the smallest attainable *p* value is 0.0625, so these tests show consistency, not significance. No result was selected on its *p* value. Sign tests and the resampling for attribution used the source model as the unit. The model panel, parameter ranges, protocols and features, the distance criteria and tolerances, the ways of building populations, the group sizes, the search-matched control and the endpoints were fixed before any selected population was analyzed. A rule fixed at the same time, which counted criteria for one explanation of the dependence against another, is not used, because the explanations are not mutually exclusive and its criteria were not independent. The Mainen model was excluded because it failed the convergence test, a decision made after the planned analyses had been run. All other analyses are exploratory: the comparisons with same-size unselected and feature-complete samples, the removal of single features, the parameter sets drawn at random, the nearest-neighbor check, the analyses without the size and passive-cable parameters and the attribution regression.

### Numerical methods

Simulations used NEURON (Carnevale and Hines, 2006) with a fixed time step for every model. A time step and spatial mesh were accepted only if all 13 features were reproduced against a reference run with a time step at least four times finer and, for models with more than one compartment, about twice as many compartments or more, on the published parameter set and five or six parameter sets drawn from the model’s own prior. A feature counted as reproduced if it lay within 5% of the reference value or within a small absolute tolerance set for each feature. Settings are listed in Table 1 and Supplementary Table S1.

## Results

### Selection can create dependence between parameters drawn independently

We first designed a test in which any dependence among parameters could come only from the way the population was built (Fig. 1A). Parameters were drawn from a prior with no dependence between them, and each selected population was compared with unselected samples and shuffled copies of the same size. The principle is easy to see in two dimensions (Fig. 1B): two independent parameters are uncorrelated (|*r*| < 0.01), but keeping only the pairs whose sum lies near a target makes them strongly negatively correlated (*r* = −0.95), although neither parameter has changed. With three or more parameters, selection can also change multivariate dependence, which we summarize with O-information.

### The Gaussian estimator detects some selection constraints and misses others

Before measuring the populations, we tested three estimators on eleven systems of known dependence (Fig. 2A). The Gaussian and Gaussian-copula estimators, which are the ones used in the studies we address, detected constraints on sums or products of positive parameters; for example, they returned −1.65 and −0.88 nats for a selection band on the sum of three log-parameters. They returned values within 0.0005 nats of zero, as for independent variables, for four systems whose dependence leaves the pattern of correlations nearly unchanged: products of zero-mean variables and selection bands on two non-monotone functions. They were therefore correct on 7 of 11 systems at every sample size, whereas the nearest-neighbor estimator, which makes no assumption about the form of the dependence, was correct on all 11 from 4,000 points upward (Fig. 2B). The Gaussian estimator thus detects some selection constraints and misses others. All results below use it unless stated otherwise, and “synergy” means synergy as this estimator measures it.

We then compared each selected population with unselected samples of exactly the same size (Fig. 3; Methods). Selection made the parameters more strongly correlated. With the root-mean-square distance, the mean absolute correlation between parameter pairs was 0.043 at a tolerance of 3 standard deviations, against 0.018 in same-size unselected samples, and 0.130 at 0.5 standard deviations, against 0.034 (Fig. 3A).

**Fig. 3.**
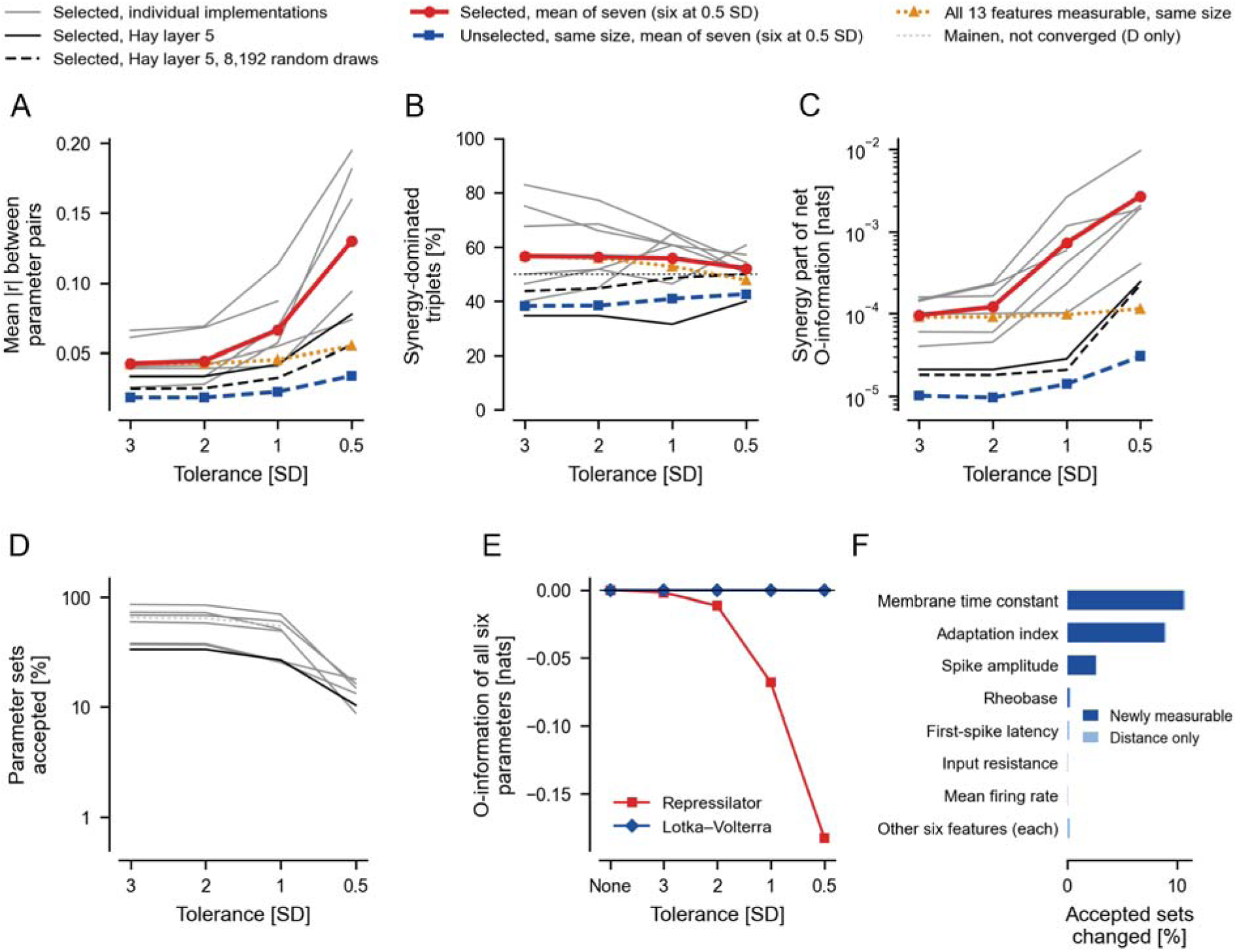
Selection for a target behavior increases dependence among parameters beyond that of unselected samples of the same size. In A–D, populations were selected by their root-mean-square distance from the published model’s 13 features, at the tolerance on the horizontal axis, in standard deviations (SD) of the unselected sample. Gray lines, individual implementations; black line, the Hay layer 5 model (main run, 1,024 parameter sets); dashed black line, the Hay model redrawn with 8,192 independent random draws (not included in the means); red line, mean of the seven converged implementations; blue dashed line, mean of 100 same-size unselected samples; orange dotted line, mean of 100 same-size samples of the unselected parameter sets whose 13 features are all measurable; all means cover six implementations at 0.5 SD, where the Migliore model retained too few sets. The Mainen model (not converged) is excluded from all means and shown only in D (dotted). Gaussian estimator throughout. A, Mean absolute Pearson correlation (|*r*|) between parameter pairs. B, Percentage of parameter triplets with negative O-information (synergy-dominated); dotted line, 50%. C, Synergy part of the net O-information (the negative part of the O-information averaged over all groups of 3 to 5 parameters, with positive values counting as zero, each divided by group size), logarithmic scale. D, Percentage of the unselected sample accepted, logarithmic scale. E, The same selection (largest-single-distance criterion) applied to two non-neuronal systems with six parameters each: a repressilator, a three-gene oscillating circuit (red squares), and a three-species Lotka–Volterra community (blue diamonds). Vertical axis, O-information of all six parameters together; “None”, the unselected sample. F, Percentage of the parameter sets accepted at 3 SD that change when one feature is left out of the acceptance rule, mean over the seven implementations; dark blue, sets that become eligible because the left-out feature was undefined for them; light blue, sets whose acceptance changes through the distance alone; bottom row, the largest value among the other six features (Supplementary Table S5). Lines end where fewer than 60 parameter sets were accepted.

### Selection raises the size of the synergy more than the share of synergistic groups

The share of synergy-dominated triplets changed little: 57% at 3 standard deviations, against 38% in same-size unselected samples, and 52% to 57% at every tolerance (Fig. 3B). With correlations this weak, the sign of a triplet’s O-information is close to that of the product of three nearly random correlations, so about 40% of triplets are negative even without selection.

The size of the synergy rose far more (Fig. 3C; Supplementary Table S3). The synergy part of the net O-information was 9.5 × 10^−5^ nats at 3 standard deviations, against 1.0 × 10^−5^ nats in same-size unselected samples, and 2.7 × 10^−3^ nats at 0.5 standard deviations, against 3.1 × 10^−5^ nats. It lay above the chance range in every model except the Hay model, whose 344 selected parameter sets exceeded it only at the strictest tolerance (Supplementary Table S4). This was a matter of sample size: redrawn with 8,192 parameter sets, the Hay model accepted 2,732 at 3 standard deviations, and their synergy part, 1.8 × 10^−5^ nats, was 19 times the mean of same-size unselected samples (chance range 0.05 to 0.16 × 10^−5^ nats; Supplementary Table S3). With parameter sets drawn at random for six models, the selected populations exceeded all 100 same-size unselected samples in all 45 combinations of model, distance criterion and tolerance (Supplementary Section S2).

Synergy grew much faster than the correlations because, for weak correlations, the O-information of a triplet is approximately the product of its three correlations (Methods). Tightening the tolerance from 3 to 0.5 standard deviations raised the mean absolute correlation about 3-fold and the synergy part about 28-fold, close to 3^3^ = 27. An exact calculation shows why selection produces synergy rather than redundancy (Methods; Supplementary Section S1): when selection narrows the population along one weighted sum of the parameters, the pairwise correlations take signs that make every triplet synergistic. A simulation of eight independent parameters matched the formula, and its synergy part rose as the 3.4th power of the mean absolute correlation. When selection narrowed two weighted sums at once, a quarter of the triplets became redundant, which is how selection on several features can raise redundancy as well. Across the 155 populations of Fig. 5C with a non-zero synergy part, the logarithm of the mean absolute correlation explained 62% of the variation in the logarithm of the synergy part. The selection effect therefore lies in the size of the O-information, which follows the strength of the correlations, not in the share of negative signs.

### Requiring measurable features and matching the target both contribute

Acceptance has two stages: all 13 features must be measurable, and their distance from the target must lie within the tolerance. To separate them, we compared each selected population with same-size samples of the unselected parameter sets whose 13 features are all measurable (Fig. 3C, orange; Supplementary Table S3). Which stage dominated at the loosest tolerance depended on the distance rule. With the root-mean-square distance at 3 standard deviations, these feature-complete samples already showed 95% of the increase in synergy over unselected samples, because requiring measurable features in practice selects models that fire repeatedly; when every feature had to lie within 3 standard deviations, they showed 37% of it. In the Hay model redrawn at random, the root-mean-square criterion at 3 standard deviations removed only 4 of 2,736 feature-complete sets, so at that tolerance its selection was the measurability requirement alone. At the strictest tolerance the match to the target dominated under both rules: the selected populations had a median of 19 times the synergy of feature-complete samples of the same size (1.8 to 57 times over six implementations), and with parameter sets drawn at random a median of 16 times (4.7 to 40 times over six). Acceptance rates are shown in Fig. 3D.

At loose tolerances, features act on selection mainly through whether they can be measured To see which features shape the acceptance boundary, we rebuilt every population selected at 3 standard deviations with each feature left out in turn (Fig. 3F; Supplementary Table S5). With the root-mean-square distance, only three features mattered: leaving out the membrane time constant, the adaptation index or the spike amplitude changed 11%, 9% and 3% of the accepted parameter sets, and almost all of these were sets that became eligible because the left-out feature had been undefined for them. Each of the other ten features changed 0.3% or less. These features therefore act through whether they can be measured, not through their values. With the largest-single-distance criterion, input resistance also acted through its value (Supplementary Section S2).

### Selection creates synergy without neuronal dynamics

The effect does not require neurons. In a repressilator, a three-gene oscillating circuit, the same selection changed the O-information of all six parameters from 2.7 × 10^−6^ nats in the unselected sample to −0.183 nats at the strictest tolerance (Fig. 3E). In a three-species Lotka–Volterra community the change had the same sign but was about 500 times smaller (−3.4 × 10^−4^ nats). Both values lay far outside the range of same-size unselected samples. The two systems differ in their dynamics, features, priors and the fraction of parameter sets retained, so how much synergy selection creates depends on the system.

### Individual synergistic groups cannot be identified reliably in small populations

Counting groups with negative O-information does not identify unusually synergistic groups: shuffled populations, which contain no dependence, already had 48% of groups negative (Fig. 4A). Groups falling beyond the search-matched surrogate threshold (Methods) were common in the five models with large selected populations (25% to 54% of groups) but rare in the two smallest (0.1% in the Hay and 1.4% in the Migliore model, with about 340 and 140 parameter sets). At the tighter tolerances, the ten most synergistic triplets also changed somewhat more than random thinning of the population to the same size would produce (Supplementary Section S2).

**Fig. 4.**
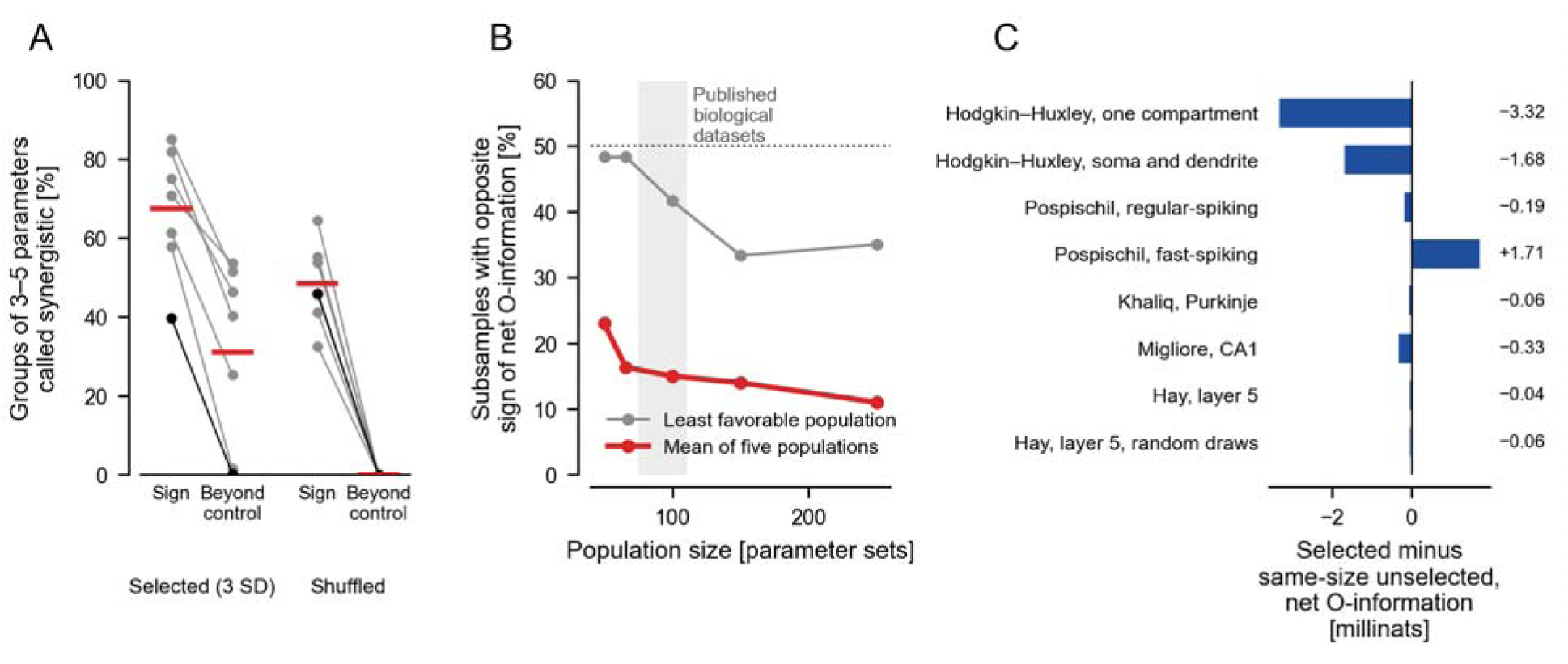
Controls for search size and population size. A, Percentage of groups of 3 to 5 parameters with negative O-information (“Sign”) and with O-information below the lower bound of the search-matched surrogate band (“Beyond control”), for populations selected at 3 standard deviations (SD) and for shuffled copies (each parameter permuted independently). Lines, implementations of the primary panel (black, Hay layer 5); red bars, means of the seven. Selected populations are averaged over both distance criteria and both protocol sets (the first set only for the Khaliq model); shuffled copies are of the populations selected with the largest-single-distance criterion. Gaussian estimator. The surrogate band uses 25 shuffled copies evaluated over the sam number of groups and is exploratory, not a calibrated family-wise error rate. B, Percentage of random subsamples whose net O-information has the opposite sign to that of the full population, as a function of subsample size; 60 subsamples per size from each of five reference populations (four implementations of the Hodgkin–Huxley and Pospischil source models). Red, mean of the five populations; gray, the least favorable population; shaded band, sizes of published biological datasets (75 to 111 cells); dotted line, 50% (a coin toss). C, Net O-information of the selected populations minus the mean of 100 unselected samples of the same size, for each implementation, averaged over tolerances, distance criteria and protocol sets (the first set only for the Khaliq model; 1 millinat = 0.001 nats); negative values indicate a shift toward synergy. Light blue, the Hay model redrawn with 8,192 random draws (fixed-amplitude steps only, both distance criteria, all tolerances). Numbers, the value of each bar. The Mainen model is excluded and not shown.

The sign of the net O-information was also unreliable at the population sizes of the biological studies we discuss, 75 and 111 cells (Fig. 4B). In subsamples of large selected populations whose own sign was stable, the sign reversed on average in 23% of subsamples of 50 parameter sets and in 11% of subsamples of 250, and in the least favorable population in 48% and 35%. At these sizes the standard error of a correlation, 0.06 to 0.14, is as large as the mean absolute correlations created by selection.

### Net O-information shifts toward synergy in six of seven models

Averaged over tolerances, distance criteria and protocol sets, selection moved the net O-information toward synergy, relative to same-size unselected samples, in six of seven implementations and four of five source models (Fig. 4C; sign test over source models, *p* = 0.375). The exception, the fast-spiking Pospischil model, moved toward redundancy (+1.7 millinats) because its redundancy rose more than its synergy; its synergy part rose, as in every model. The shift toward redundancy came from the populations selected with the largest-single-distance criterion at 1 and 0.5 standard deviations, mainly the two at 0.5 with only 66 and 86 parameter sets (+15.1 and +13.0 millinats); all eight populations selected with the root-mean-square distance moved toward synergy, including the two in Supplementary Table S3.

The Hay model’s average shift was small in both runs (−0.04 millinats in the main run and −0.06 with 8,192 random draws; Fig. 4C), because its synergy and redundancy parts are both small, but with the random draws all seven of its selected populations lay below every same-size unselected sample: at 3 and 0.5 standard deviations its net O-information was −1.5 and −17.6 × 10^−5^ nats, against chance ranges of −0.08 to +0.09 and −0.6 to +0.7 × 10^−5^ nats (Supplementary Table S3).

Selection raised the redundancy part as well (Supplementary Table S3). At 0.5 standard deviations it lay above the chance range in all six implementations with a population at that tolerance, counting the Hay model drawn at random (in the main run, its 106 sets did not separate from chance), but it rose less than the synergy part. In the selected populations the synergy part was a median of 74 times the redundancy part at 3 standard deviations and 3.6 times at 0.5, against about 1 in same-size unselected samples. Reva and colleagues (2025) reported synergy about ten times stronger than redundancy. Their summary differs from ours, so the two ratios cannot be compared directly, but selection alone produced ratios of the same order or larger.

### Selection also creates non-linear pairwise dependence

Because the Gaussian estimator sees only correlations, we applied the nearest-neighbor estimator to the 27 selected populations large enough for it, all from parameter sets drawn at random (2,002 to 7,101 sets from six models; Methods). In the 23 populations from the five models other than the Hay model, the mutual information between parameter pairs exceeded that of synthetic populations with the same rank correlations, by 0.002 to 0.015 nats per pair, and 17 of the 23 exceeded all ten same-size unselected samples compared in the same way (Fig. 2D; Supplementary Table S6). In the four Hay populations it did not (−0.003 to −0.001 nats per pair), and none exceeded the unselected samples. Selection therefore also creates pairwise dependence that the correlations miss, in five of the six models tested.

Averaged over all triplets, the nearest-neighbor O-information showed no consistent departure from the synthetic populations (−0.008 to +0.009 nats; Fig. 2C), a range similar to that of unselected samples compared with their own synthetic populations. The Gaussian shift on the same triplets, 0.05 to 2.3 millinats (under 0.01 millinats in the Hay model), lies well within this spread, so the nearest-neighbor estimator could neither confirm nor rule it out.

### Parameters that set cell size carry much of the synergy in two models

Membrane capacitance, diameter and, in the Migliore model, axial resistance set how conductance densities translate into currents and voltages, so a feature that depends on a conductance and the membrane area constrains them jointly. To see how much of the synergy these size and passive-cable parameters carry, we recomputed it with them left out of the parameter vector (Supplementary Section S2, Table S7). At 0.5 standard deviations the synergy part fell from 206 to 23 × 10^−5^ nats in the regular-spiking and from 191 to 49 × 10^−5^ nats in the fast-spiking Pospischil model, but stayed far above the chance range (at most 2.0 × 10^−5^ nats). In both, the most synergistic triplet was the leak conductance, the leak reversal potential and the soma diameter, and triplets containing a size parameter carried 78% to 92% of the synergy, although they were 64% to 71% of all triplets. In the two Hodgkin–Huxley models the synergy part rose when these parameters were left out, and in the Khaliq model it changed little (41 to 32 × 10^−5^ nats). In the Migliore model, with 127 to 154 selected sets, it fell inside the chance range at every tolerance. The parameter sets drawn at random gave the same pattern. Leaving parameters out of the measure is not the same as holding them fixed in the simulations, which we did not do.

### How a population is built changes the balance between synergy and redundancy

We next built populations in other ways (Fig. 5A). The share of synergy-dominated triplets was 40% in same-size unselected samples, 11% when neighboring parameters were correlated in the prior, 62% in a clustered population built without any selection for behavior and 60% in populations selected at 3 standard deviations; the other constructions lay between. The clustered population shows the limit of the Gaussian reading: this estimator scored it as synergistic, whereas the nearest-neighbor estimator, which sees the clusters, scored it as strongly redundant (0.21 to 0.49 nats per triplet above its shuffled copies).

**Fig. 5.**
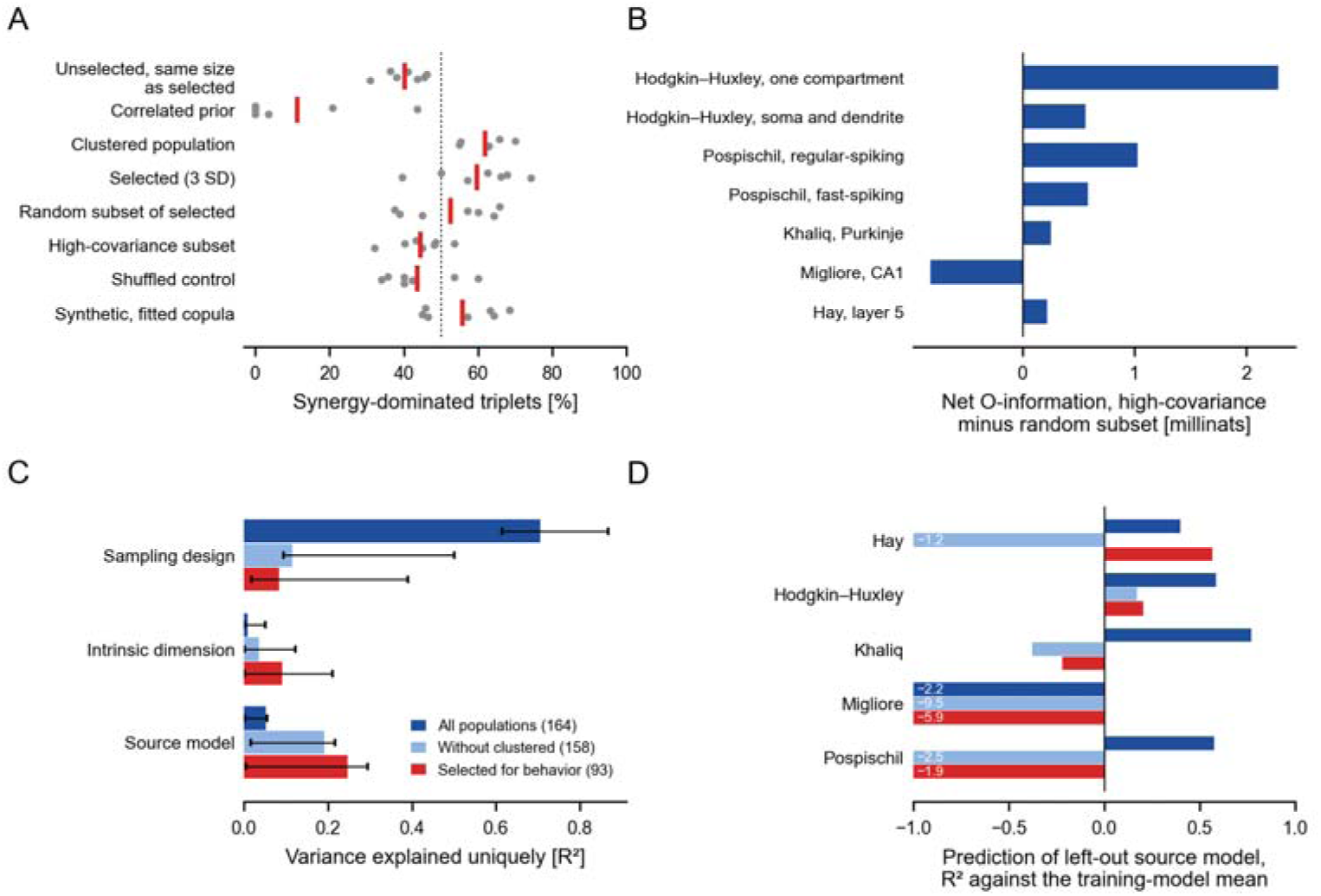
Population construction and the attribution of synergy. A, Percentage of synergy-dominated parameter triplets for eight ways of building a population from the same models. Gray dots, implementations of the primary panel; red bars, means; dotted line, 50%. Selected populations use the largest-single-distance criterion at 3 standard deviations (SD). The unselected samples, shuffled copies and synthetic populations have the size of the selected populations; the random and high-covariance subsets have half that size; the correlated prior and clustered population were simulated separately (six implementations). Population sizes therefore differ between rows, and the share of synergy-dominated triplets depends on size (Fig. 3B). B, Net O-information of the high-covariance subset minus that of the size-and mean-matched random subset, for each implementation (1 millinat = 0.001 nats); positive values indicate a shift toward redundancy. C, Variance in the synergy part of the net O-information explained uniquely by each group of predictors (the R² lost when that group is removed from an ordinary least-squares regression; standardized outcome; construction method, tolerance and source model as fixed indicator variables), for all 164 populations (dark blue; full regression R² = 0.83), without the six clustered populations (light blue; 0.59) and for the 93 populations selected for behavior (red; 0.70). Population geometry is represented by intrinsic dimension only, because measures computed from the covariance matrix are functions of the same correlations as the synergy part. Error bars, 95% intervals from 2,000 resamples of source models. D, R² for predicting the synergy of each source model’s populations from sampling design and intrinsic dimension, with that source model left out of the fit, relative to predicting the training models’ mean (positive values beat that prediction), for the same three sets of populations; values below −1 are truncated and printed.

With the target behavior fixed, choosing the half of a selected population with the strongest covariance shifted the net O-information toward redundancy, relative to a random subset matched in size and parameter means, in six of seven models (Fig. 5B). Reva and colleagues (2025) found the same shift with a different panel, code and selection criterion. In parameter sets drawn at random, the nearest-neighbor estimator agreed in five of six models.

### Construction and source model cannot be separated among selected populations

As a descriptive analysis, we asked how the variation in synergy across populations relates to how they were built and to their source model (Fig. 5C). Over all 164 populations, sampling design explained 0.71 of the variation uniquely and source model 0.05 (Supplementary Table S2). This depended on the six clustered populations, whose synergy part was about 100 times that of the selected populations. Without them, the unique shares were 0.12 for sampling design and 0.19 for source model, and among the 93 populations selected for behavior 0.08 and 0.25, with overlapping intervals.

Predicting each source model from the others gave the same picture (Fig. 5D). Predictions from sampling design and intrinsic dimension beat the training models’ mean for four of five source models over all populations but for two of five among the selected populations. Against the left-out model’s own mean, they did so for four of five and none of five. The fit therefore did not reliably predict the synergy of selected populations from a new source model, and among these populations this panel cannot separate construction from source model.

## Discussion

Here, we show that selecting parameter sets drawn from a factorized prior by the model’s own electrical output is sufficient, without any co-regulation of parameters, to produce synergy of the kind reported in populations of neuron models, as the usual estimator measures it. Compared with unselected samples of the same size, the synergy part increased in all seven converged models, also when six of them were redrawn at random, and the net O-information shifted toward synergy in six of seven. Selection also created non-linear pairwise dependence in five of the six models tested (Fig. 2D). A population of acceptable models is therefore not only a collection of plausible solutions but a distribution shaped by the operations that generated and filtered it. These results measure the selection effect in model populations; they are not a re-analysis of any published biological dataset and do not show that reported co-regulation is absent.

A key result is that the synergy statistic responds similarly across systems sharing only the procedure. Selection made independent parameters synergistic in a single-compartment squid-axon model, cerebellar and hippocampal models, and a gene circuit (Fig. 3), because the acceptance rule constrains functions of all parameters. Approximately additive constraints, such as a feature depending on a sum or a product of conductances, make the selected parameters negatively correlated, and the Gaussian estimator scores a triplet with three negative correlations as synergistic (Fig. 2A); for a single such constraint, every triplet becomes synergistic (Supplementary Section S1). Part of the constraint is simply that the features can be measured, which selects models that fire repeatedly and have a measurable membrane time constant (Figs. 3C, F); this stage operates in any pipeline that rejects models with undefined features. But selection acts through each model’s mapping from parameters to features, explaining why its effect varied and was about 500 times smaller in the Lotka–Volterra community than in the repressilator. Thus, the statistic captures the joint influence of model and acceptance rule and cannot distinguish them alone.

This account predicts a large effect when the features used for selection are nearly additive functions of several parameters’ logarithms, the prior is broad relative to the accepted region, and the accepted population is large relative to the parameter groups examined. Effects should be hard to detect when few parameter sets are available: with 344 sets the Hay model barely separated from chance, but with 2,732 it did so clearly. The roughly one-half share of synergy-dominated triplets should not be generalized; chance alone produces a similar share at these sizes.

### Relation to previous work

These results extend prior findings. Reva and colleagues (2025) proposed stochastically sampled model populations as a null model for biological variability; our results measure what such a null contains once the population is selected for behavior. They also found that covariance-only model selection, with targets and structure unchanged, shifted results toward redundancy, suggesting that interaction structure reflects population-level statistical organization. We found the same shift in six of seven independently coded models, compared with size-and parameter-mean-matched random subsets, despite using a different panel and selection criterion, although with the same covariance-based estimator. Their column-shuffling control addressed search size, not selection. Their estimator, like ours, reads synergy from the pattern of pairwise correlations (Fig. 2A). Schneider et al. (2023) showed that functional constraints induce negative correlations among independently generated variables, largely independent of correlations in the generating population. This is the same phenomenon: in our selection band on the sum of three log-parameters, the three negative correlations are what the Gaussian estimator scores as strong synergy. In epidemiology, selection on a common effect can induce positive or negative associations depending on how causes combine in the selection probability (Cole et al., 2010). Acceptance criteria are such rules, which may explain why effect sizes differed across models and between the two non-neuronal systems.

Parameter interactions also arise in model dynamics without selection. In the Hodgkin–Huxley cable model, propagating uncertainty in original rate-constant fits through hundreds of thousands of simulations yielded small first-order but large total-order Sobol indices (Sobol’, 2001) during spiking, indicating that excitability depends mainly on parameter interactions (Korngreen, 2026a). These findings complement, rather than contradict, ours: variance-based sensitivity analysis examines how parameters combine to produce model output, whereas a selected population’s O-information measures how parameter values depend on one another after fixing the output. The former reflects model equations under a specified prior; the latter combines the equations with the acceptance rule. Selection can induce dependence without non-additive dynamics: even an additive mapping from independent parameters to a shared output can yield dependence when sets are selected for that output. This does not show that parameters interact dynamically, so we use “synergy” only statistically.

Our results do not contradict population-independent biological co-regulation. Channel gene expression in identified neurons correlates in cell-type-specific patterns and changes with neuromodulation (Schulz et al., 2007; Temporal et al., 2012); homeostatic tuning also produces correlations among conductances measured in identified neurons (O’Leary et al., 2013). Co-regulation can coexist with selection-induced dependence. We question only using O-information computed in a selected population as evidence of co-regulation without a control that reproduces selection. For biological datasets, this control must model how cells were included and characterized. If selection or classification used electrical criteria, such as firing pattern or input resistance, sample a cell-type model from a factorized prior, apply the same criteria, measure the same features in the same number of cells, and use the same estimator. Dependence beyond this reference would show that the model, prior, and selection do not fully explain the data, but would not establish biological co-regulation. Conversely, matching the reference would not show that co-regulation is absent. In mouse cardiac myocytes, a compensation between calcium and potassium conductances, predicted by models selected for a normal calcium transient, was validated with currents measured in genetically diverse mouse strains (Rees et al., 2018).

This raises a methodological point about documenting and reusing model ensembles. We proposed releasing reusable neuron models as ensembles with a declared scope, explicit uncertainty, and an assembly record (Korngreen, 2026b). Our results show that this record must include the prior and acceptance rule: ensembles of the same neuron built with different tolerances, distance criteria, or covariance-based subsetting have different higher-order structure (Figs. 3 and 5). Without these details, readers cannot determine which structure belongs to the neuron. The same applies to simulation-based inference posteriors, whose dependence structure depends on the prior, summary features, and simulator (Gonçalves et al., 2020; Tolley et al., 2024).

### Limitations

Several limitations apply. The first concerns the information measure. We used the Gaussian estimator because it is the one used in the studies we address, but it depends only on the correlation matrix: a negative value summarizes pairwise structure and does not prove dependence beyond it, the estimator misses constraints that leave the correlations unchanged (Fig. 2A), and agreement between its Gaussian and Gaussian-copula versions is not independent confirmation. The test systems had effects above 0.5 nats, so classifying them correctly does not show sensitivity to the millinat-sized changes measured in the neuron models. The nearest-neighbor estimator was usable only on parameter sets drawn at random, for the six models redrawn that way, because it responds to the spacing of points in an evenly spread design (Supplementary Sections S1 and S2). Its average over triplets can hide departures of opposite sign and cannot separate pairwise from three-way dependence, so whether selection creates dependence among three parameters beyond their pairwise structure remains untested, and these measurements do not bound the synergy another estimator might report.

Second, the populations. The priors are expert-chosen ranges, not measured conductance distributions, and say nothing about biological variability. Five independent source models, with no neuron class represented twice, cannot establish rules for neuron classes or separate source-model identity from architecture, number of parameters and implementation history; the shares of variation attributed to construction and to source model depended on which populations were included (Fig. 5C). The unselected sample came from an evenly spread quasi-random design, which has less chance dependence than independent random draws; the main comparison was repeated with random draws for six models with the same result. Against shuffled copies, which behave like independent draws, the results held for the five models with large populations but, except at the strictest tolerance, not for the two with fewer than 350 parameter sets (Supplementary Table S4); redrawn with 8,192 parameter sets, the Hay model showed the effect at every tolerance, whereas the Migliore model, with 256, was not tested at a larger size. The size and passive-cable parameters were left out of the measure, not held fixed in the simulations, so how much synergy remains when they cannot vary is not known. The estimates of sign reliability come from four implementations of two source models; they motivate resampling but do not estimate the uncertainty of any biological dataset, whose measurement noise, heterogeneity and inclusion criteria differ. The high-covariance subset matched the random subset in size and parameter means but not in each parameter’s full distribution, so it shows that selecting by covariance changes the balance of synergy and redundancy, not that covariance alone causes the change. The Mainen model failed the convergence test and was excluded; including it did not alter any qualitative conclusion (Supplementary Section S1).

Third, the features. Rheobase was measured with a 140 ms step and a 0 mV spike criterion, whereas the other spike features used 400 ms steps and a −20 mV criterion. Leaving rheobase out of the acceptance rule at 3 and 2 standard deviations changed on average 0.3% of accepted parameter sets and the synergy part by 2%, so it does not explain the results, although this does not bound the effect of a differently measured rheobase (Supplementary Section S1). The spike width used is the width at the −20 mV detection threshold, 0.1 to 0.3 ms wider than the width at half amplitude in the published parameter sets. Firing output entered the distance twice, as spike count and as firing rate; removing the rate changed the strictest populations by about 9% of their members and did not change the conclusions (Supplementary Section S2). In the Migliore model the resting potential did not vary across the unselected sample, so it did not enter the distance. All features were measured at the soma, so the dendritic properties that the Hay and Migliore models were built to reproduce, such as back-propagating action potentials and dendritic calcium spikes, did not constrain selection, and selection on dendritic features could create different dependence. Tolerances were expressed in standard deviations of the unselected sample, not in the measured variability of each feature, which would define a different accepted region.

Finally, the interpretation. We quantified dependence created by selection but did not test it against a population with biological co-regulation, nor did we test whether parameters identified as synergistic interact in the dynamics of the model. These results therefore neither show that higher-order structure in real neurons is an artifact nor rule out co-regulated groups of channels.

### Recommendations for analyzing model populations

Claims about parameter dependence in a model population require a reproducible account of its construction: parameter prior, simulator, stimulation protocols, measured features, acceptance criterion, tolerance and population size. Five practices strengthen statistical interpretations.

First, use a selection-matched reference: sample a factorized prior, then apply the same model and acceptance procedure. Rebuild this control for each model, feature set and tolerance, as effect sizes varied several-fold across our models.

Second, compare each statistic with unselected or shuffled samples of equal size, and report O-information magnitude, not just the share of groups with a negative sign.

Third, compare individual groups against a search-matched control evaluated across the same number of groups.

Fourth, report population size, acceptance rate, the share of models rejected because a feature could not be measured, and the share of resamples preserving the statistic’s sign.

Fifth, pair the Gaussian estimator with at least one estimator that makes no Gaussian-dependence assumption, and compare it with shuffled copies and with synthetic populations that have the same correlations, not only with the Gaussian estimate. Do not apply nearest-neighbor estimators to populations drawn from evenly spread (quasi-random) designs: selection keeps the design’s spacing, and the estimator responds to it.

These controls answer different questions: the selection-matched reference estimates the dependence induced by the prior, model and acceptance rule, whereas the equal-size and search-matched comparisons assess finite-sample and multiple-search effects. Together they help determine whether an inference about a selected population’s internal structure exceeds plausible procedural explanations. Passing them does not establish biological co-regulation or functional interaction. To claim that parameters interact dynamically, jointly perturb them in the model along population-occupied directions, and resolve the interaction by order.

## Conclusion

Selecting model populations for a shared behavior is enough, without any co-regulation, to create a synergy-dominated statistic of the kind reported for neuron models, as the usual estimator measures it. How large and how synergistic that dependence appears reflects the prior, the selection rule, the size of the search, the population size and the estimator, as well as the modeled neuron. Treating neuron models as ensembles rather than single parameter sets is an important step, but the ensemble is itself shaped by its construction, and inferences from its parameter structure must be evaluated against the prior, selection procedure and matched controls that define it.

## Supporting information

Supplemental results

## Author Contribution

Alon Korngreen: conceptualization, methodology, software, formal analysis, writing – original draft, writing – review and editing. Nurit Degani-Katzav: conceptualization, investigation, writing – review and editing.

## Statement about AI use

Claude Code was used extensively in developing the simulation and statistical analysis software, implementing validation procedures, and maintaining the computational workflow. Claude was also used to assist with manuscript editing and revision. The authors directed the scientific investigation, evaluated the computational results, made the methodological and interpretive decisions, and take responsibility for the accuracy of the analyses and the conclusions.

## Data Availability

All code, analysis tables and figures are available at https://github.com/alon67/COMPASS (release tag v0.4.11-manuscript-revision). Model source files are not redistributed, because their licenses permit use but not redistribution; the repository’s fetch script downloads them from

ModelDB and checks each file against a recorded checksum, so every result can be traced to the exact model code. The scripts that draw the figures and tables read only the release’s stored analysis tables and reproduce them exactly. Scripts that recompute analyses from individual simulations read the per-simulation feature tables, which are not deposited because of their size (several gigabytes) but are regenerated from the code and the recorded random seeds. No experimental data were generated for this study.

## Ethics Statement

No human or animal subjects were used in this study.

