## Supplemental results for "Selection for a shared behavior creates parameter dependence read as synergy in populations of neuron models"

Section S1 summarizes the numerical and analysis settings and gives details of analysis choices that bear on the main results and the exact result for a single selection constraint. Section S2 gives, model by model, the effect of selection on synergy, redundancy and net O-information, the comparison with unselected samples of the same size, the removal of single features from the acceptance rule, the nearest-neighbor check on parameter sets drawn at random, the stability of the most synergistic triplets and the analysis without the parameters that set cell size and passive cable properties.

### S1. Analysis details

#### Summary of numerical and analysis settings

**Table S1. Summary of numerical and analysis settings.**

| Item | Setting |
|---|---|
| Simulator | NEURON 8.2.6, driven from Python 3.12.4; each model run in its own process |
| Integration | Fixed time step (main-text Table 1); adaptive integration not used |
| Convergence test | All 13 features reproduced (within 5% or a small absolute tolerance) against a reference time step of 0.00625 ms or finer and, for multi-compartment models, a refined mesh; published parameter set plus 5 prior draws (6 for the Migliore and Mainen models) |
| Reproducibility test | 3,341 stored parameter sets of the main runs, at least one from each of their 3,057 stored blocks, and 1,893 of the random-draw run, at least one from each of its 1,804 blocks, were simulated again and reproduced every stored feature (absolute difference at most $10^{-6}$ in the feature's own units, $10^{-9}$ nA for rheobase, or $10^{-9}$ times the stored value, whichever is larger, to allow for reruns on other computers); all stored parameter values were regenerated from their recorded random seeds |
| Parameter sampling | Factorized log-uniform prior (no dependence imposed between parameters); scrambled Sobol low-discrepancy design (pseudo-random draws in a repeat run for six models); ¼ to 4 times the published value for conductances |
| Simulations | 38,400 parameter sets from the factorized prior across eight models; 18,944 from the correlated prior and 18,944 from the clustered design (six models); 76,288 in the main run; 45,056 drawn at random for six models; 121,344 in all |
| Protocols | Rest 300 ms; one hyperpolarizing and two depolarizing 400 ms steps after 120 ms delay; rheobase search (64-point scan plus bisection, 140 ms step, spike = upward crossing of 0 mV) |
| Feature extraction | eFEL 5.7.29 for spike amplitude, width at −20 mV and after-hyperpolarization depth; firing features from upward crossings of −20 mV; both checked against an independent implementation |
| Selection | All features measurable; distance to target in standard deviations of the unselected sample; largest single distance or root-mean-square (weighted Euclidean) distance; tolerance 3, 2, 1 or 0.5; at least 60 accepted sets |
| O-information | Gaussian and Gaussian-copula estimators with small-sample correction; groups of 3, 4 and 5 parameters; all groups if ≤1,200, otherwise a fixed random sample of 1,200 |
| Same-size comparison | 100 random subsamples of the unselected sample, 100 of its feature-complete sets and 20 shuffled copies per selected population, each the size of that population |

| Item | Setting |
|---|---|
| Search-matched control | 25 shuffled copies per population, same number of groups; band from the 5th percentile of the minima to the 95th percentile of the maxima |
| Feature removal | Each of 13 features left out of the acceptance rule in turn; tolerances 3 and 2 SD; both distance criteria; newly eligible sets separated from changes in distance |
| Estimator test | 11 known systems (7 four-variable; 4 built from positive log-uniform parameters) × 4 sample sizes (300 to 16,000) × 3 estimators (Gaussian, Gaussian copula, nearest-neighbor); classification threshold ±0.02 nats |
| Nearest-neighbor check | All triplets and pairs of 27 selected populations drawn at random (2,002 to 7,101 sets), rank-transformed to normal scores; 10 each of unselected samples of the same size, shuffled copies and fitted Gaussian-copula populations; fourth nearest neighbor |
| Sign reliability | 60 subsamples at each of 5 sizes (50 to 250) from 5 reference populations (four implementations, two source models) |
| Attribution | Ordinary least squares; unique $R^2$; 2,000 resamples of source models; all populations, without clustered populations, selected populations only |
| Random seeds | Sampling seeds recorded for each simulation run in the repository's records; analysis seed 20260804, used for group sampling and controls |
| Software | NumPy 1.26.4, SciPy 1.14.0, pandas 2.2.3, statsmodels 0.14.6, scikit-learn 1.5.2 |

#### Unit of analysis

The resampling for attribution, the cross-validation folds and the sign tests use the source model rather than the implementation, because implementations of the same source model share equations and parameter conventions and are not independent. Subsampling within a population and shuffling of parameter columns use other units by design. Where the number of units is small, intervals are described as descriptive rather than as calibrated confidence statements.

#### Attribution of variation across models

The regression uses three groups of predictors: sampling design, population geometry and source model. Implementation and model architecture (number of parameters, number of compartments, and whether the model has a single compartment) are not included. Each implementation belongs to exactly one source model, and the architecture variables are fixed for each implementation, so the implementation indicators contain all the information in the source-model indicators and in the architecture variables. With all of them in the regression, removing source model, implementation or architecture would leave the fitted values unchanged, and the unique variance of each would be zero by construction rather than by measurement.

Because source model, architecture and implementation history are not independently varied, this analysis describes associations across the sampled populations, not the causal contribution of each model characteristic. Population geometry is represented only by the two intrinsic-dimension estimates. The four other geometry measures (mean absolute correlation, effective rank, participation ratio and the share of variance on the first principal component) are computed from the covariance matrix, and the synergy part is itself a function of the correlation matrix, so they explain it partly by construction: with them, geometry alone explained 71% of the variation, and the unique variances were 0.155 for sampling design, 0.053 for geometry and 0.008 for source model.

The two Khaliq populations selected with the second protocol set are excluded, because that set is undefined for a model whose published parameter set has a rheobase of zero, leaving 164 populations. With sampling design and the intrinsic-dimension estimates, the prediction for a left-out source model did not beat the training-mean prediction for the Migliore model ($R^2$ = −2.18); sampling design alone beat it for all five ($R^2$ 0.17 to 0.77). With the covariance-based geometry measures added, prediction beat the training-mean prediction for four of five source models in all three sets of populations (Migliore −3.8 over all populations), but those measures explain the synergy part partly by construction. These scores compare the prediction with predicting the training models' mean for every left-out population. Scored against the left-out model's own mean, which a prediction beats only if it is accurate in absolute value within that model, the counts of positive scores over all populations, without the clustered populations and among the selected populations were 4, 1 and 0 of 5 with the covariance-based geometry measures, 4, 0 and 0 with intrinsic dimension and 3, 0 and 0 with sampling design alone (for example, the Migliore prediction from sampling design alone over all populations scored +0.62 against the training mean and −3.4 against its own mean).

Both results depend on the six clustered populations: without them, the unique variances were 0.115 for sampling design, 0.035 for intrinsic dimension and 0.191 for source model ($R^2$ = 0.586), and for the 93 populations selected for behavior 0.084, 0.091 and 0.247 ($R^2$ = 0.699); predictions from sampling design, alone or with intrinsic dimension, beat the training-mean prediction for at most two of five source models in either set. Splitting sampling design into choices made in advance (construction method and tolerance) and outcomes that depend on the model (acceptance rate and population size) gave unique variances of 0.687 and 0.017 over all populations. Bootstrap draws containing a single source model were skipped (5 of 2,000).

**Table S2. Attribution of variation in the synergy summary across 164 populations (five source models, seven implementations).**

| Group of predictors | Variance explained by the group alone ($R^2$) | Variance explained only by this group (unique $R^2$) | 95% interval over source models |
|---|---|---|---|
| Sampling design (construction method, tolerance, population size, acceptance rate) | 0.762 | 0.706 | 0.614 to 0.868 |
| Population geometry (intrinsic dimension, estimated in two ways) | 0.006 | 0.009 | 0.0015 to 0.050 |

| Group of predictors | Variance explained by the group alone ($R^2$) | Variance explained only by this group (unique $R^2$) | 95% interval over source models |
|---|---|---|---|
| Source model | 0.088 | 0.052 | 0.0003 to 0.054 |

All three groups together explain $R^2$ = 0.834. Unique $R^2$ is the $R^2$ of the full regression minus that of the regression without the group. Intervals: 2.5th and 97.5th percentiles over 2,000 resamples of source models with replacement.

We do not form a ratio between unique variances: the unique variance of source model came close to zero in some resamples, so such a ratio has no useful upper bound. With five source models and no neuron class represented by two of them, source model, architecture, number of parameters and implementation history cannot be separated, and the small unique share of source model is as consistent with that limitation as with source model being unimportant. The Hay populations are analyzed with the two parameters that change no output removed, as in every other analysis.

#### Rheobase definition

Rheobase, one of the 13 selection features, was measured with a 140 ms step and counted a spike as an upward crossing of 0 mV, whereas all other spike features were measured during 400 ms steps with spikes detected at −20 mV. The two definitions disagree in practice. In the stored simulations, nine recordings at 1.5 times rheobase had their first spike later than 140 ms (five in the Hay model, three in the one-compartment and one in the two-compartment Hodgkin–Huxley model; the latest at 280.8 ms), and 2,453 of 67,078 spiking recordings at the stronger fixed-amplitude step (3.7%) had spike peaks at or below 0 mV, 1,402 of them in the Khaliq model. To bound the effect of this inconsistency we rebuilt every population selected at 3 and 2 standard deviations with rheobase left out of the acceptance rule altogether (Section S2, Table S5). With the root-mean-square distance at 3 standard deviations this changed 0.3% of the accepted parameter sets on average (at most 0.7%, in the Migliore model) and changed the synergy part of the net O-information by 1.8% on average (at most 11%); with the largest-single-distance criterion it changed 0.2% of accepted sets. Leaving rheobase out therefore does not explain the results at the tolerances tested (3 and 2 standard deviations); it does not show what a differently measured rheobase would do, which could change which parameter sets are measurable and, in the second protocol set, the injected current and every spike feature. The populations analyzed in the main text were selected with fixed-amplitude steps, so rheobase enters them as one feature of 13; the second protocol set, which scales the step to each parameter set's own rheobase, depends on it more strongly; it enters only the pooled summaries of main-text Figs. 4A, 4C and 5C, D, and it is undefined for the Khaliq model, whose published parameter set fires spontaneously (rheobase zero).

**Resolution of the rheobase search**

The rheobase search scanned 64 equally spaced amplitudes before refining. This guarantees that any range of currents that evokes spikes and is wider than 1/64 of the search range is found; a narrower range can be missed.

**High-covariance subset**

The high-covariance subset was chosen within bands of distance to the target and matched in size and parameter means to the random subset, but not in the full distribution of each parameter. Across the 72 parameters of the eight implementations the mean Kolmogorov–Smirnov distance between the two subsets was 0.064 (maximum 0.18), 18% of comparisons differed at nominal $p < 0.05$, and the ratio of standard deviations ranged from 0.79 to 1.28. The comparison in main-text Fig. 5B therefore shows that selecting by covariance changes the balance of synergy and redundancy, not that covariance alone causes the change.

**Parameters that change nothing**

Every parameter was checked for whether scaling it fourfold up or down changes at least one measured output. Two parameters of the Hay model (apical Kv3-like potassium and apical low-voltage-activated calcium conductance) did not; three somatic conductances of a replication model not analyzed in the main text also did not. These parameters were removed before populations were built.

**Choice of estimator**

The main analyses use Gaussian and Gaussian-copula estimators, which depend only on the correlations: main-text Fig. 2 shows that they detect constraints on sums or products of positive parameters but are blind, at every sample size tested, to dependence that leaves the correlations unchanged. They were chosen because they are the estimators used in the literature under discussion. The nearest-neighbor estimator was applied to parameter sets drawn at random for the six models redrawn that way (Section S2, Table S6). It found non-linear pairwise dependence created by selection in five of them, but not in the Hay model; the mean triplet O-information showed no consistent departure from synthetic populations with the same correlations, which does not bound dependence among three parameters. On the evenly spread Sobol design the estimator responds to the spacing of the points, so it was applied only to parameter sets drawn at random (Section S2). In the smaller populations the effect of a non-Gaussian estimator remains unknown. The Gaussian and Gaussian-copula estimates correlated at 0.91 across 14,291 triplet estimates in the 164 populations of the primary panel (each unselected population counted once; 0.89 when every stored copy is counted), and their shares of synergy-dominated triplets correlated at 0.96.

**Synergy created by a single selection constraint**

Let the logarithms of n parameter factors be independent and standardized, $x \sim N(0, I)$, and let selection keep the draws whose weighted sum $z = u \cdot x$, with $|u| = 1$, lies in a band. The components of x orthogonal to u are independent of z, so the kept draws have covariance $I - k\, uu^T$, where $k = 1 - \mathrm{Var}(z \mid \text{band})$ lies between 0 and 1; for the band $|z| < \tau$, $k = 2\tau\varphi(\tau)/(2\Phi(\tau) - 1)$, where $\varphi$ and $\Phi$ are the standard normal density and distribution function. For a triplet, write $a_i = u_i^2$ and $s = a_1 + a_2 + a_3$. Every principal submatrix of $I - k\, uu^T$ has determinant $1 - k$ times the sum of its $a_i$, so the Gaussian O-information of the triplet is exactly $\Omega = ½[\ln(1 - ks) + \Sigma_i \ln(1 - ka_i) - \Sigma_i \ln(1 - k(s - a_i))]$. Expanding the logarithms in k, the first- and second-order terms cancel, and the leading term is $\Omega \approx -k^3 a_1 a_2 a_3$. The correlations are $r_{ij} = -ku_iu_j/\sqrt{[(1 - ka_i)(1 - ka_j)]} \approx -ku_iu_j$, so the leading term is the product of the three correlations, and it is negative whatever the signs of the weights. Evaluated on a grid of k from 0.001 to 0.999 for 200 random weight vectors, the exact $\Omega$ was never positive: a single constraint of this kind makes every triplet synergistic, and its synergy grows as the cube of the correlations it creates.

We checked the result numerically. We drew 400,000 sets of eight independent standard normal parameters, kept those whose projection on a random unit vector lay within $\pm\tau$, for $\tau = 3$, 2, 1, 0.5 and 0.25 standard deviations, and computed the Gaussian O-information of all 56 triplets. The estimates matched the exact formula to within 0.0013 nats, and at $\tau = 0.5$ the leading term correlated with the exact value at 0.98. Over the five bands, the synergy part rose as the 3.4th power of the mean absolute correlation, close to the cube expected from the leading term. With two constraints along orthogonal directions (both bands ±0.5), 75% of the triplets were synergistic and 25% redundant, because two constraints no longer impose one sign pattern on the correlations. Selection on 13 features constrains several combinations of parameters at once, and the redundancy part of the selected populations rose as well (Table S3). The calculation assumes Gaussian parameters and a linear constraint, whereas the model parameters are log-uniform and the features are non-linear functions of them, so it explains the direction of the effect, not its size in any model.

**Exclusion of the Mainen model**

At its production settings (time step 0.0125 ms, 2,304 compartments) the Mainen model reproduced 12 of the 13 selection features on all seven test parameter sets (the published set and six drawn from the prior) against a reference refined in both time (0.00625 ms) and space (4,440 compartments). The adaptation index failed on one test set: a spike close to threshold was lost from a nine-spike train, and this ratio of interspike intervals changed from −0.045 to +0.068. The failure is one of spatial convergence in one feature, not a numerical instability. A coarser mesh (1,066 compartments) failed six features, so the refinement used was necessary; a finer one would roughly double the cost of the most expensive model in the panel and would not remove spikes close to threshold. It was therefore excluded from all summaries, because a model that fails a numerical test is excluded from primary inference rather than footnoted. The decision depended only on the convergence test, which involves no information measure. With the Mainen model included, selection moved the net O-information toward synergy in five of six source models (exact two-sided sign test, $p = 0.22$), against four of five without it, and the high-covariance subset moved toward redundancy in four of six; no qualitative conclusion changed. In the Mainen model itself, populations selected at 3 standard deviations had 37% to 39% synergy-dominated triplets and negative net O-information with the weighted Euclidean distance.

**The full unselected sample is not a fair reference**

In the full unselected sample only 2.7% of triplets were synergy-dominated, against 56% in the selected populations, but this comparison overstates the effect of selection. The full sample was larger than any selected population and was drawn from a scrambled Sobol sequence, which spreads points more evenly than independent random draws. Its mean absolute correlation between parameters was at most 0.0015 in six of the seven models of the primary panel, whereas independent random draws of the same sizes (256 to 8,192 parameter sets) would give 0.009 to 0.05. With correlations this small, the small-sample correction of the Gaussian estimator made almost every group appear slightly redundant. The exception shows the mechanism: in the Khaliq model, 595 of 4,096 parameter sets were removed as numerical failures (main text, Methods), so its unselected sample is less evenly spread, and 19% of its triplets were synergy-dominated, against 0% in the other six models. The main text therefore compares selected populations with equal-sized unselected subsamples and shuffled copies (Section S2), while distinguishing the low-discrepancy reference from the approximately independent-draw behavior of the shuffled reference.

### S2. Results for each model

These analyses used the stored simulations of the main run, except for the comparisons on parameter sets drawn at random, which used 45,056 additional simulations (121,344 in all with the 76,288 of the main run). They cover the primary panel (seven implementations of five source models, with the two Hay parameters that change no output removed) and use the Gaussian estimator unless stated otherwise.

#### Comparison with unselected samples and shuffled copies of the same size

For every selected population we drew 100 random subsamples of the unselected sample and 20 shuffled copies of the selected population itself, each of exactly the same size, and analyzed them identically (Tables S3 and S4). The share of synergy-dominated triplets in the selected population lay within the range of the unselected samples in four of seven models; the synergy part of the net O-information lay above that range in six of seven, and above the range of the shuffled copies in the five models with large populations.

The comparison was repeated on parameter sets drawn with independent pseudo-random draws instead of the Sobol design, for the five models with the largest selected populations (36,864 parameter sets, the same sizes as the main run). At 3 standard deviations the synergy part was $1.2 \times 10^{-4}$ nats on average, against $5.1 \times 10^{-6}$ nats in unselected samples of the same size, and it lay above all 100 such samples in all 38 analyzable combinations of model, distance criterion and tolerance; the mean absolute correlation was 0.045 against 0.015, and the share of synergy-dominated triplets 56% against 46%. The Hay model, whose 1,024 Sobol draws gave only 344 sets at 3 standard deviations, was redrawn with 8,192 random draws: all seven of its analyzable selected populations (604 to 2,732 sets) lay above all 100 same-size unselected samples, with a synergy part of $1.8 \times 10^{-5}$ nats at 3 standard deviations against $0.10 \times 10^{-5}$ nats.

A second reference separates the two stages of acceptance: 100 samples of the same size drawn from the unselected parameter sets whose 13 features are all measurable. The features most often undefined were the adaptation index and the first interspike interval, which need repeated spiking at the stronger step, the spike amplitude and the membrane time constant. At least three spikes is necessary but not sufficient: in the single-compartment Hodgkin–Huxley model 4,364 unselected sets fired at least three times and 3,065 had all 13 features (Pospischil fast-spiking, 7,233 and 7,208); between 34% (Hay, 344 of 1,024) and 88% (Pospischil fast-spiking) of the unselected sets had all 13. At 3 standard deviations the distance criterion removed at most 1.2% of the feature-complete sets (none in the Hay model), and the selected populations exceeded all 100 feature-complete samples in only two of six models (Hodgkin–Huxley soma and dendrite, Pospischil fast-spiking); at 0.5 standard deviations they exceeded them in five of six models, and in all six with parameters drawn at random. Over all analyzable combinations of model, distance criterion, protocol set and tolerance, the selected population exceeded all 100 feature-complete samples in 54 of 90 on the Sobol design (excluding the two Khaliq populations of the undefined second protocol set) and in 33 of 45 with parameters drawn at random. With firing rate removed from the distance (it equals 2.5 times spike count), the populations at 0.5 standard deviations changed by 9% of their members on average, their synergy part changed by −1% (Sobol design) and −10% (random draws), and they still exceeded all 100 feature-complete samples in five of six and five of five models.

These are complementary reference distributions rather than interchangeable null samples: unselected subsamples retain the original quasi-random design, whereas shuffling retains the selected marginals but destroys cross-parameter dependence.

**Table S3. Effect of selection in each implementation.**

| Tolerance (SD) | Sets accepted | Synergy part ($10^{-5}$ nats) | | | Redundancy part ($10^{-5}$ nats) | | Net O-information ($10^{-5}$ nats) | |
|---|---|---|---|---|---|---|---|---|
| | | Selected | Same-size unselected: mean (95% range) | Same-size feature-complete | Selected | Same-size unselected: mean (95% range) | Selected | Same-size unselected: 95% range |
| **Hodgkin–Huxley, one compartment** | | | | | | | | |
| 3 | 3,055 | 14.7 | 0.04 (0.002 to 0.15) | 14.7 | 0.73 | 0.04 (0.005 to 0.13) | −14.0 | −0.12 to 0.12 |
| 0.5 | 1,091 | 952 | 0.39 (0.02 to 1.60) | 16.8 | 16.3 | 0.35 (0.02 to 1.29) | −936 | −1.54 to 1.16 |
| **Hodgkin–Huxley, soma and dendrite** | | | | | | | | |
| 3 | 3,130 | 15.8 | 0.05 (0.006 to 0.13) | 14.8 | 0.22 | 0.04 (0.007 to 0.09) | −15.6 | −0.10 to 0.06 |
| 0.5 | 1,467 | 184 | 0.19 (0.04 to 0.53) | 15.6 | 0.60 | 0.19 (0.06 to 0.50) | −183 | −0.28 to 0.29 |
| **Pospischil, regular-spiking** | | | | | | | | |
| 3 | 6,054 | 5.99 | 0.004 (0 to 0.01) | 5.98 | 0.005 | 0.005 (0.001 to 0.01) | −5.98 | −0.007 to 0.009 |
| 0.5 | 717 | 206 | 0.57 (0.09 to 1.47) | 7.62 | 80.2 | 0.64 (0.18 to 1.50) | −125 | −1.01 to 1.08 |
| **Pospischil, fast-spiking** | | | | | | | | |
| 3 | 7,119 | 3.99 | 0.001 (0 to 0.003) | 3.68 | 0.001 | 0.001 (0.001 to 0.004) | −3.99 | −0.003 to 0.003 |
| 0.5 | 1,224 | 191 | 0.26 (0.02 to 0.77) | 4.50 | 93.3 | 0.28 (0.03 to 0.79) | −97.9 | −0.68 to 0.61 |
| **Khaliq, Purkinje** | | | | | | | | |
| 3 | 2,432 | 9.89 | 3.62 (2.94 to 4.56) | 9.77 | 0.13 | 0.06 (0.02 to 0.13) | −9.76 | −4.50 to −2.91 |
| 0.5 | 572 | 40.6 | 4.69 (1.53 to 8.03) | 10.7 | 26.1 | 1.31 (0.49 to 2.87) | −14.5 | −7.33 to 0.14 |
| **Migliore, CA1** | | | | | | | | |
| 3 | 154 | 14.2 | 2.07 (0.10 to 7.37) | 12.5 | 2.05 | 2.97 (1.20 to 5.92) | −12.1 | −5.75 to 5.26 |
| 0.5 | fewer than 60 | — | — | — | — | — | — | — |
| **Hay, layer 5: main run (1,024 parameter sets)** | | | | | | | | |
| 3 | 344 | 2.13 | 1.36 (0.46 to 2.50) | 2.13 | 1.28 | 1.32 (0.70 to 2.24) | −0.84 | −1.34 to 1.29 |
| 0.5 | 106 | 24.6 | 12.3 (5.51 to 21.7) | 13.3 | 8.03 | 12.4 (7.79 to 20.8) | −16.5 | −9.81 to 8.85 |
| **Hay, layer 5: 8,192 parameter sets drawn at random** | | | | | | | | |
| 3 | 2,732 | 1.83 | 0.10 (0.05 to 0.16) | 1.81 | 0.33 | 0.10 (0.05 to 0.19) | −1.49 | −0.08 to 0.09 |

| Tolerance (SD) | Sets accepted | Synergy part ($10^{-5}$ nats) | | | Redundancy part ($10^{-5}$ nats) | | Net O-information ($10^{-5}$ nats) | |
|---|---|---|---|---|---|---|---|---|
| | | Selected | Same-size unselected: mean (95% range) | Same-size feature-complete | Selected | Same-size unselected: mean (95% range) | Selected | Same-size unselected: 95% range |
| 0.5 | 792 | 22.5 | 0.64 (0.30 to 1.17) | 2.77 | 4.96 | 0.63 (0.28 to 1.25) | −17.6 | −0.61 to 0.67 |

Populations selected with the root-mean-square distance and fixed-amplitude steps, at 3 and 0.5 standard deviations (SD) of the unselected sample. Synergy and redundancy parts: the negative and positive parts of the O-information averaged over groups of 3 to 5 parameters, each divided by group size (main-text Methods); the net O-information is the redundancy part minus the synergy part, so negative values indicate synergy. Same-size samples: 100 samples, each the size of the selected population, drawn from all unselected parameter sets or from those whose 13 features are all measurable (feature-complete); their central 95% is the chance range. The Hay model is shown for the main run and for 8,192 parameter sets drawn independently at random. —, fewer than 60 parameter sets accepted. The change in net O-information averaged over all selection conditions is shown in main-text Fig. 4C.

#### Table S4. Populations selected at 3 standard deviations (root-mean-square distance) compared with unselected samples and shuffled copies of the same size.

| Implementation | Parameter sets | Synergy-dominated triplets, selected [%] | Same, unselected samples [%] (95% range) | Same, shuffled copies [%] | Synergy part, selected [nats] | Synergy part, unselected samples [nats] | Synergy part, shuffled copies [nats] |
|---|---|---|---|---|---|---|---|
| Hodgkin–Huxley, one compartment | 3,055 | 40 | 43 (22 to 63) | 52 | $1.5 \times 10^{-4}$ | $4.0 \times 10^{-7}$ (up to $1.5 \times 10^{-6}$) | $1.0 \times 10^{-6}$ (up to $2.9 \times 10^{-6}$) |
| Hodgkin–Huxley, soma and dendrite | 3,130 | 50 | 44 (29 to 61) | 46 | $1.6 \times 10^{-4}$ | $4.5 \times 10^{-7}$ (up to $1.3 \times 10^{-6}$) | $9.5 \times 10^{-7}$ (up to $2.8 \times 10^{-6}$) |
| Pospischil, regular-spiking | 6,054 | 75 | 38 (22 to 54) | 45 | $6.0 \times 10^{-5}$ | $3.9 \times 10^{-8}$ (up to $1.1 \times 10^{-7}$) | $3.2 \times 10^{-7}$ (up to $8.0 \times 10^{-7}$) |
| Pospischil, fast-spiking | 7,119 | 83 | 30 (11 to 50) | 48 | $4.0 \times 10^{-5}$ | $9.0 \times 10^{-9}$ (up to $3.4 \times 10^{-8}$) | $2.2 \times 10^{-7}$ (up to $5.0 \times 10^{-7}$) |
| Khaliq, Purkinje | 2,432 | 68 | 43 (32 to 52) | 45 | $9.9 \times 10^{-5}$ | $3.6 \times 10^{-5}$ (up to $4.6 \times 10^{-5}$) | $1.2 \times 10^{-6}$ (up to $2.5 \times 10^{-6}$) |
| Migliore, CA1 | 154 | 46 | 30 (12 to 50) | 42 | $1.4 \times 10^{-4}$ | $2.1 \times 10^{-5}$ (up to $7.4 \times 10^{-5}$) | $1.1 \times 10^{-4}$ (up to $2.6 \times 10^{-4}$) |
| Hay, layer 5 | 344 | 35 | 39 (30 to 47) | 41 | $2.1 \times 10^{-5}$ | $1.4 \times 10^{-5}$ (up to $2.5 \times 10^{-5}$) | $2.1 \times 10^{-5}$ (up to $3.0 \times 10^{-5}$) |

Synergy part: the mean size of the negative O-information values across groups of 3 to 5 parameters, each divided by group size (main text, Methods). "Up to": the 97.5th percentile over 100 unselected samples or 20 shuffled copies. The unselected samples come from an evenly spread quasi-random design and are therefore quieter than independent random draws; the shuffled copies behave like independent draws.

### Removing one feature from the acceptance rule

#### Table S5. Effect of leaving one feature out of the acceptance rule at 3 standard deviations.

| Feature left out | Accepted sets changed, root-mean-square distance [%] | Change in synergy part [%] (range over models) | Models with negative net O-information | Accepted sets changed, largest distance [%] | Change in synergy part [%] (range over models) | Models with negative net O-information |
|---|---|---|---|---|---|---|
| Resting potential | 0.1 | +1 (-1 to +5) | 7 of 7 | 0.0 | -0 (-0 to +0) | 7 of 7 |
| Input resistance | 0.0 | +0 (-1 to +2) | 7 of 7 | 2.7 | -28 (-80 to +5) | 7 of 7 |
| Membrane time constant | 10.7 | +0 (-34 to +48) | 7 of 7 | 10.5 | -3 (-23 to +23) | 6 of 7 |
| Sag ratio | 0.2 | -2 (-14 to +0) | 7 of 7 | 0.6 | +2 (+0 to +11) | 7 of 7 |
| Rheobase | 0.3 | +2 (-1 to +11) | 7 of 7 | 0.2 | -0 (-3 to +0) | 7 of 7 |
| First-spike latency | 0.2 | +0 (-1 to +5) | 7 of 7 | 1.8 | -2 (-6 to +0) | 7 of 7 |
| Spike amplitude | 2.6 | -11 (-51 to +10) | 6 of 7 | 2.4 | -6 (-24 to +5) | 6 of 7 |
| Spike width | 0.1 | +1 (-1 to +5) | 7 of 7 | 0.1 | -1 (-6 to +1) | 7 of 7 |
| After-hyperpolarization depth | 0.1 | +1 (-1 to +5) | 7 of 7 | 0.2 | -1 (-5 to +0) | 7 of 7 |

| Feature left out | Accepted sets changed, root-mean-square distance [%] | Change in synergy part [%] (range over models) | Models with negative net O-information | Accepted sets changed, largest distance [%] | Change in synergy part [%] (range over models) | Models with negative net O-information |
|---|---|---|---|---|---|---|
| First interspike interval | 0.2 | +1 (-1 to +5) | 7 of 7 | 0.4 | +2 (-4 to +20) | 7 of 7 |
| Mean firing rate | 0.0 | -0 (-1 to +0) | 7 of 7 | 0.0 | +0 (+0 to +0) | 7 of 7 |
| Adaptation index | 9.0 | -18 (-65 to -1) | 6 of 7 | 7.8 | -14 (-39 to +3) | 6 of 7 |
| Spike count | 0.0 | -0 (-1 to +0) | 7 of 7 | 0.0 | +0 (+0 to +0) | 7 of 7 |

Means over the seven implementations of the primary panel. With all 13 features, net O-information was negative in all seven models with the root-mean-square distance and in six of seven with the largest-single-distance criterion. Because features can constrain the same parameter combinations, these one-at-a-time omissions measure the effect of removing each feature conditional on the other twelve, not its independent contribution to selection. Almost all of the changes with the root-mean-square distance were sets that became eligible because the left-out feature had been undefined for them (membrane time constant 10.6 of 10.7 percentage points, adaptation index 8.8 of 9.0, spike amplitude 2.6 of 2.6); with the feature-complete population held fixed, leaving out the adaptation index or the spike amplitude changed the synergy part by 0.6% and 0.1% on average. With the largest-single-distance criterion, the 28% reduction for input resistance involved no newly eligible sets.

### Nearest-neighbor check on parameter sets drawn at random

#### Table S6. Nearest-neighbor O-information of parameter triplets in populations selected from parameter sets drawn at random, compared with three references.

| Implementation | Parameter sets | Triplets | Selected minus unselected samples of the same size [nats] (range) | Selected minus shuffled copies [nats] (range) | Selected minus synthetic populations with the same rank correlations [nats] (range) |
|---|---|---|---|---|---|
| Hodgkin–Huxley, one compartment | 3,007 | 20 | +0.005 (−0.007 to +0.014) | +0.001 (−0.017 to +0.013) | +0.002 (−0.006 to +0.015) |
| Hodgkin–Huxley, soma and dendrite | 3,186 | 56 | −0.005 (−0.013 to +0.000) | −0.009 (−0.021 to +0.004) | −0.008 (−0.018 to +0.001) |
| Pospischil, regular-spiking | 6,098 | 56 | −0.001 (−0.006 to +0.004) | +0.003 (−0.004 to +0.011) | +0.002 (−0.003 to +0.006) |
| Pospischil, fast-spiking | 7,101 | 35 | +0.001 (−0.003 to +0.009) | +0.000 (−0.010 to +0.008) | +0.000 (−0.008 to +0.006) |
| Khaliq, Purkinje | 2,433 | 120 | +0.002 (−0.004 to +0.007) | −0.002 (−0.012 to +0.012) | +0.000 (−0.007 to +0.009) |
| Hay, layer 5 | 2,732 | 286 | −0.002 (−0.007 to +0.002) | −0.004 (−0.010 to +0.003) | −0.006 (−0.012 to −0.001) |

Populations selected at 3 standard deviations with the root-mean-square distance from parameter sets drawn at random. Each parameter was converted to normal scores by rank before estimation; fourth nearest neighbor. Values are the mean over all triplets of the selected population minus that of each of ten references; ranges over the ten. The synthetic populations have the selected population's rank correlations and parameter distributions (Gaussian copula), so the last column reflects any dependence the correlations miss, within pairs or among three parameters; as a mean over triplets it can hide departures of opposite sign (in the Khaliq population, individual triplets differed by −0.078 to +0.076 nats). Calibration: the full unselected samples differed from their own shuffled copies by −0.005 to +0.003 nats and from their own synthetic populations by −0.006 to +0.007 nats. Over all 27 selected populations with at least 2,000 parameter sets, from six models (both distance criteria; populations at 3, 2 and 1 standard deviations, none at 0.5), the differences ranged from −0.007 to +0.013 nats against unselected samples and from −0.008 to +0.009 nats against synthetic populations, whereas the Gaussian shift on the same triplets was 0.05 to 2.3 millinats toward synergy (0.04 to 2.2 millinats on the same normal scores) in the five models other than the Hay model, and under 0.01 millinats in the Hay model. Pairwise nearest-neighbor mutual information exceeded that of the synthetic populations in all 23 populations of those five models, by 0.002 to 0.015 nats per pair, but not in the four Hay populations (−0.003 to −0.001 nats) (full unselected samples, −0.001 to +0.003 nats). Calibrated against 10 unselected samples of the same size, each compared with five synthetic populations of its own (−0.006 to +0.005 nats), 17 of the 27 selected populations exceeded all ten, none of them from the Hay model; with the estimator of Kraskov et al. (2004) and 39 synthetic populations, the excess at 3 standard deviations was 0.002 to 0.011 nats per pair, with 32 to 39 of the 39 below the selected value, in those five models, and −0.002 nats in the Hay model, with 6 of the 39 below it.

Why the Sobol design could not be used: a selected population contains every Sobol point inside a region and so keeps the design's even spacing, whereas a random subsample, a shuffled copy or a synthetic population does not, and the nearest-neighbor estimator responds to spacing. On the Sobol design the selected populations of this table were 0.011 to 0.072 nats per triplet below unselected samples of the same size, but a box-shaped region of the same design with the same number of points, which creates no dependence, was 0.010 to 0.082 nats below them (Hodgkin–Huxley one compartment −0.082, soma and dendrite −0.060, Pospischil regular-spiking −0.010, fast-spiking −0.011, Khaliq −0.015), and against shuffled copies the selected populations were 0.24 to 0.42 nats lower. The selected populations differed from the boxes by −0.023 to +0.025 nats, so the control shows that the comparison is sensitive to spacing, not that spacing explains all of the shift; in the Khaliq model the boxes were drawn after 595 of 4,096 sets had been removed by the voltage check. The box control, its parameters and its per-triplet results are deposited with the code.

#### Stability of the most synergistic triplets

The list holds up to ten triplets, those with negative O-information (eight in the single-compartment Hodgkin–Huxley population at 3 standard deviations). Of the triplets on the list at 3 standard deviations (root-mean-square distance), on average 9.3 remained on the list at 2 standard deviations, 6.3 at 1 and 3.7 at 0.5 standard deviations; with the largest-single-distance criterion the values were 6.6, 3.5 and 5.0. With the root-mean-square distance, random subsets of the 3-standard-deviation population, reduced to the size of each tighter population (200 per implementation and tolerance), kept 9.3, 7.4 and 4.9. The strong overlap between loose tolerances therefore reflects the near-identity of those populations, not independent evidence that the groups are reliably identified; the tighter populations turned over somewhat more of the list than thinning alone would.

#### Leaving out the parameters that set cell size and passive cable properties

We recomputed the Gaussian summaries of every selected population of the main run and of the random-draw run, and of the same 100 same-size unselected samples, with membrane capacitance and diameter (capacitance and axial resistance in the Migliore model) left out of the parameter vector (Table S7). The Hay model varies none of these parameters. The synergy part stayed above all 100 unselected samples in 75 of the 76 populations of the Hodgkin–Huxley, Pospischil and Khaliq models (both runs, both distance criteria, all tolerances) and fell inside their range in all five Migliore populations. In the Khaliq model the synergy of the unselected samples ($3.6 \times 10^{-5}$ nats at 3 standard deviations) came almost entirely from these parameters ($0.10 \times 10^{-5}$ nats without them). In the populations selected at 3 and 0.5 standard deviations, triplets that contain a size or cable parameter carried 78% to 92% of the order-3 synergy in the Pospischil models (64% to 71% of all triplets), 53% to 64% in the Khaliq model (53%), 91% in the Migliore model (64%) and 2% to 33% in the Hodgkin–Huxley models (64% to 80%), whose most synergistic triplets combined the somatic sodium and potassium conductances. In both

Pospischil models the most synergistic triplet was the leak conductance, the leak reversal potential and the soma diameter. The five most synergistic triplets of each population, with their correlations, are deposited with the code.

**Table S7. Synergy part with and without the parameters that set cell size and passive cable properties.**

| Implementation | Parameters left out | Tolerance (SD) | Sets | All parameters: selected | All parameters: unselected, mean (95% range) | Left out: selected | Left out: unselected, mean (95% range) |
|---|---|---|---|---|---|---|---|
| Hodgkin–Huxley, one compartment | capacitance, soma diameter | 3 | 3,055 | 14.7 | 0.04 (0.002 to 0.15) | 45.0 | 0.03 (0 to 0.19) |
| Hodgkin–Huxley, one compartment | capacitance, soma diameter | 0.5 | 1,091 | 952 | 0.39 (0.02 to 1.60) | 1317 | 0.25 (0 to 1.99) |
| Hodgkin–Huxley, soma and dendrite | capacitance, dendrite diameter | 3 | 3,130 | 15.8 | 0.05 (0.006 to 0.13) | 42.7 | 0.04 (0.001 to 0.19) |
| Hodgkin–Huxley, soma and dendrite | capacitance, dendrite diameter | 0.5 | 1,467 | 184 | 0.19 (0.04 to 0.53) | 337 | 0.19 (0.007 to 0.76) |
| Pospischil, regular-spiking | capacitance, soma diameter | 3 | 6,054 | 5.99 | 0.004 (0 to 0.01) | 3.67 | 0.004 (0 to 0.02) |
| Pospischil, regular-spiking | capacitance, soma diameter | 0.5 | 717 | 206 | 0.57 (0.09 to 1.47) | 23.3 | 0.56 (0.02 to 2.00) |
| Pospischil, fast-spiking | capacitance, soma diameter | 3 | 7,119 | 3.99 | 0.001 (0 to 0.003) | 2.14 | 0.001 (0 to 0.006) |
| Pospischil, fast-spiking | capacitance, soma diameter | 0.5 | 1,224 | 191 | 0.26 (0.02 to 0.77) | 49.3 | 0.36 (0.001 to 1.64) |
| Khaliq, Purkinje | capacitance, soma diameter | 3 | 2,432 | 9.89 | 3.62 (2.94 to 4.56) | 10.1 | 0.10 (0.006 to 0.34) |
| Khaliq, Purkinje | capacitance, soma diameter | 0.5 | 572 | 40.6 | 4.69 (1.53 to 8.03) | 32.2 | 1.33 (0.15 to 3.12) |
| Migliore, CA1 | capacitance, axial resistance | 3 | 154 | 14.2 | 2.07 (0.10 to 7.37) | 1.07 | 2.03 (0.02 to 10.5) |
| Migliore, CA1 | capacitance, axial resistance | 1 | 127 | 58.8 | 3.81 (0.30 to 13.9) | 18.3 | 3.91 (0.009 to 20.7) |

Populations of the main run selected with the root-mean-square distance and fixed-amplitude steps; the Migliore model had no population at 0.5 standard deviations (SD), so its population at 1 SD is shown. Synergy part in $10^{-5}$ nats, Gaussian estimator. Unselected: 100 random samples of the unselected parameter sets, each the size of the selected population, analyzed with the same parameters; 95% range, their central 95%. Leaving parameters out of the measure is not the same as holding them fixed in the simulations.
